# Diacylglycerol at the inner nuclear membrane fuels nuclear envelope expansion in closed mitosis

**DOI:** 10.1101/2022.06.01.494365

**Authors:** Sherman Foo, Amaury Cazenave-Gassiot, Markus R. Wenk, Snezhana Oliferenko

## Abstract

Nuclear envelope (NE) expansion must be controlled to maintain nuclear shape and function. The nuclear membrane expands massively during ‘closed’ mitosis, enabling chromosome segregation within an intact NE. Phosphatidic acid (PA) and diacylglycerol (DG) can both serve as biosynthetic precursors for membrane lipid synthesis. How they are regulated in time and space and what are the implications of changes in their flux for mitotic fidelity is largely unknown. Using genetically encoded PA and DG probes, we show that DG is depleted from the inner nuclear membrane during mitosis in the fission yeast *Schizosaccharomyces pombe*, but PA does not accumulate, indicating that it is rerouted to membrane synthesis. We demonstrate that DG-to-PA conversion catalysed by the diacylglycerol kinase Dgk1 and direct glycerophospholipid synthesis from DG by diacylglycerol cholinephosphotransferase / ethanolaminephosphotransferase Ept1 reinforce NE expansion. We conclude that DG consumption through both *de novo* and the Kennedy pathways fuels a spike in glycerophospholipid biosynthesis, controlling NE expansion, and ultimately, mitotic fidelity.

## Introduction

The double-membrane nuclear envelope (NE) is a hallmark of eukaryotic cells. It consists of the outer nuclear membrane (ONM) continuous with the ER, and the inner nuclear membrane (INM) facing the nucleoplasm. The NE is perforated by the nuclear pore complexes that regulate the transport and exchange of macromolecules between the cytoplasm and the nucleus. Tight control of nuclear shape and size is essential for proper cell function, and aberrant regulation of nuclear morphology has been implicated in aging and disease (Webster et al., 2009; Cantwell and Dey, 2021). The spherical shape of the interphase nucleus minimizes the amount of membrane material required to make the NE. Changes in lipid biosynthesis, nuclear trafficking and/or INM-chromatin attachments may all lead to deviation from this simplest shape (Webster et al., 2009; Arnone et al., 2013; Cantwell and Dey, 2021).

‘Closed’ mitosis of lower eukaryotes, in which the nucleocytoplasmic integrity is maintained throughout chromosome partitioning, represents a fascinating example of highly regulated NE expansion that is not scaled to nuclear volume increase (Zhang and Oliferenko, 2013). The model fission yeast *Schizosaccharomyces pombe* undergoes ‘closed’ mitosis, whereas its relative, *Schizosaccharomyces japonicus* breaks and reforms the NE (Yam et al., 2011). We have previously linked this divergence to differences in nuclear membrane management. As nuclear volume remains constant in ‘closed’ mitosis, the surface area of the mother nucleus must increase to allow for the formation of two daughter nuclei. The nuclear membrane expands dramatically in *S. pombe* but not *S. japonicus*, necessitating NE breakdown in this organism (Yam et al., 2011). Massive expansion of the NE during mitosis in *S. pombe* is driven by the CDK1-dependent inactivation of an evolutionarily conserved regulator of phosphatidic acid (PA) flux, lipin. This does not occur in *S. japonicus*, likely due to differences in trans-regulation of lipin phosphorylation (Makarova et al., 2016). Lipin inactivation upon mitotic entry presumably causes a block in the conversion of PA to diacylglycerol (DG) and a diversion of PA towards production of glycerophospholipids (GPL), thus triggering NE expansion required for ‘closed’ nuclear division (Makarova et al., 2016).

The subcellular distribution of the lipin substrate PA and its product DG at the NE and the possible implications of changes in their distribution for NE expansion and mitotic fidelity remain unknown. Current models of how the regulation of PA flux controls NE growth are largely based on lipidomics analyses of extracts from cells with constitutively (e.g., gene deletion or mutation) or slowly (e.g., overexpression) modified PA regulation network (Santos-Rosa et al., 2005; Han et al., 2008). Yet, in such cases cells settle at metabolic steady states different from the wild-type, limiting the insight into dynamic events occurring in mitosis.

In the budding yeast, which also undergoes ‘closed’ mitosis, lipin deficiency leads to PA accumulation (Han et al., 2008). It was proposed that such an increase in PA might lead to changes in the biophysical properties of the nuclear membrane (Han et al., 2008). This, together with PA-dependent transcriptional upregulation of glycerophospholipid biosynthetic genes (Loewen et al., 2004) could lead to increased nuclear membrane biogenesis. Yet, this regulation circuitry is not conserved in fission yeasts (Rhind et al., 2011; Rutherford et al., 2022); it is also not clear if lipin inactivation leads to increased PA levels in *S. pombe*. To gain an insight into the mechanisms underlying mitotic NE expansion it is imperative to distinguish between different subcellular pools of PA and DG and probe their fates in normal mitosis and upon deregulation of the relevant biosynthetic pathways.

Here we address these questions by using a combination of imaging, genetic and cell physiological approaches in *S. pombe* and the related fission yeast *S. japonicus*.

## Results

### Development of genetically encoded biosensors to probe subcellular distribution of PA and DG in *S. pombe*

We have built a set of genetically encoded PA and DG biosensors, extending the designs used to visualize these lipid classes in *Saccharomyces cerevisiae* (Romanauska and Kohler, 2018). Our PA and DG sensors were expressed from the medium-strength *S. pombe cdc15* promoter and tagged at the C-terminus with a GST biochemical tag and a fluorophore (Fig. 1A). Appending or omitting SV40 NLS sequences around the lipid-binding module has allowed us to probe lipid enrichment in the nucleus or the cytoplasm.

**Figure 1.**
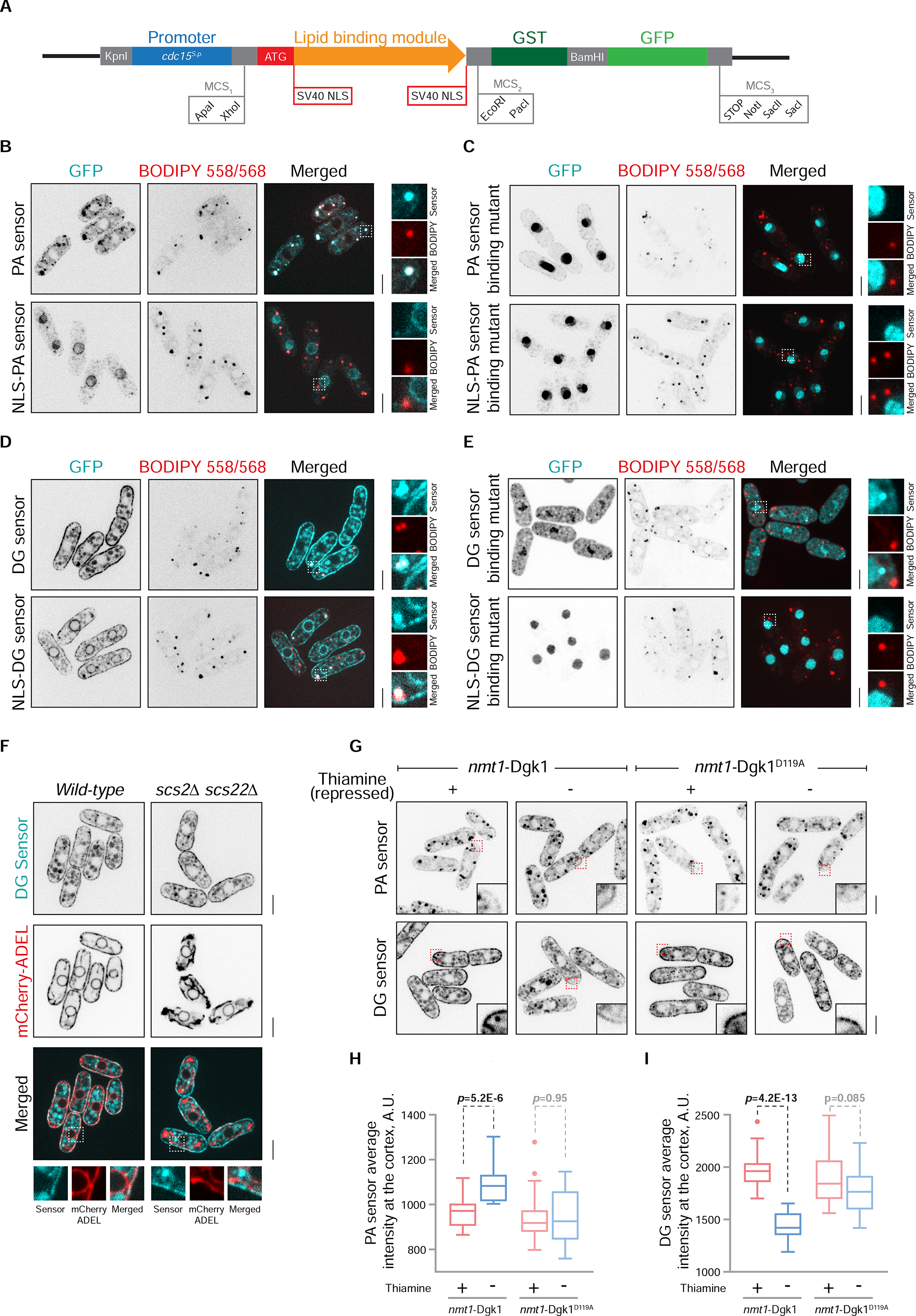
Lipid biosensors for probing subcellular distribution of PA and DG in fission yeast. (**A**) A diagram depicting the design of lipid sensors used in this study. The lipid sensors consist of a lipid-binding module (*S. cerevisiae* Opi1 PA-binding amphipathic helix and *R. norvegicus* PKCβ DG-binding domain) under the control of the *S. pombe cdc15* promoter, with a C-terminal GST-tag and a fluorophore. In the NLS versions of the sensors, the lipid-binding module is flanked by two SV40 NLS motifs. (**B**) Single-plane spinning-disk confocal images of *S. pombe* expressing the PA and NLS-PA sensors show PA enrichment at LDs and the INM, respectively. LDs are stained with BODIPY 558/568. (**C**) Mutations in the PA-binding domain abolish normal localization of the PA sensors. Note that a weak bipartite NLS appearing in the mutated PA-binding domain results in its nuclear re-localization in the absence of the SV40 NLS. (**D**) Single-plane spinning-disk confocal images of *S. pombe* cells expressing the DG and NLS-DG sensors show DG enrichment at the cortex, LDs and the INM. (**E**) Mutations predicted to abolish DG-binding lead to re-localization of the sensors. The DG-binding mutant sensor becomes largely cytoplasmic whereas the NLS version is enriched in the nucleoplasm. (**F**) DG sensor localizes to the cortex in cells where the ER (marked by mCherry-ADEL) is largely detached from the plasma membrane due to the absence of the VAP proteins Scs2 and Scs22. (**G**) Overexpression of the DG kinase Dgk1 increases PA and decreases DG at the cortex. No significant changes are observed upon overexpression of the catalytically inactive D119A mutant of Dgk1. The wild-type and the mutant Dgk1 proteins are expressed under the control of the thiamine-repressible *nmt1* promoter. Quantitations of these phenotypes are shown in (**H**) and (**I**), n=20 cells, p-values are derived from the unpaired t-test. In panels (**B**-**G**), insets represent magnified areas indicated by the dotted lines; scale bars represent 5µm.

The PA recognition module was engineered from the Q2 domain of the *S. cerevisiae* transcriptional factor Opi1 (Loewen et al., 2004), using further optimization enhancing its PA-binding properties (Hofbauer et al., 2018). Specifically, the PA sensor used throughout this study was constructed by removing the endogenous NLS within the codon-optimized Q2 helix, introducing a G120W point mutation predicted to increase PA-binding selectivity, and duplicating the helix to stabilize it at the membrane (Fig. S1A, B). In exponentially growing *S. pombe*, the cytoplasmic PA sensor was enriched mildly at the cellular cortex and strongly in distinct puncta, confirmed to be lipid droplets (LDs), as visualized by staining cells with the neutral lipid dye BODIPY 558/568 (Fig. 1B, upper panel). Using the NLS version of the sensor, we detected PA enrichment at the INM (Fig. 1B, lower panel). Confirming sensor specificity, mutation of amino acid residues known to mediate PA binding (Loewen et al., 2004) abrogated the specific localization patterns (Fig. 1C). Note that these mutations yield a weak bipartite NLS and result in nuclear import even in the case of the cytoplasmic construct.

The DG biosensor was developed based on the duplicated codon-optimized *Rattus norvegicus* protein kinase C (PKCβ C1a/b) DG-binding domain (Lucic et al., 2016) (Fig. 1D and Fig. S1C, D). DG was strongly enriched in the INM and the cellular cortex, as well as in some but not all LDs (Fig. 1D). The DG-binding specificity was validated by using DG-binding deficient versions of the sensor containing two point mutations (Q63E in the C1a domain and Q128E in the C1b domain) (Lucic et al., 2016). The loss of DG binding resulted in the redistribution of constructs to the nucleoplasm and/or cytoplasm depending on the presence of the NLS (Fig. 1E). The cortical localization of the DG sensor remained unaffected in the absence of ER-plasma membrane attachments in *scs2*Δ *scs22*Δ genetic background (Zhang et al., 2012), indicating that DG is enriched at the plasma membrane but not the peripheral ER in *S. pombe* (Fig. 1F).

We validated both sensors further by analysing their distribution under conditions where relative abundance of PA and DG was expected to deviate from the wild-type. Overexpression of the diacylglycerol kinase Dgk1, which catalyses the reverse reaction to lipin by phosphorylating DG to produce PA, from the strong thiamine-repressible *nmt1* promoter resulted in an increase in PA sensor signal at the cellular cortex (Fig. 1G, H), with a concomitant decrease in cortical DG (Fig. 1G, I). These phenotypes were not observed in cells overexpressing the D119A catalytically inactive mutant of Dgk1 (Han et al., 2008).

The phosphatase Spo7-Nem1 positively regulates lipin activity, and the loss of either subunit of the phosphatase complex phenocopies the lack of lipin (Siniossoglou et al., 1998; Santos-Rosa et al., 2005; Makarova et al., 2016). Liquid chromatography electrospray ionization tandem mass spectrometry analyses of total cellular extracts of *S. pombe* lipin pathway mutants revealed a slight decrease in PA levels in the *spo7*Δ mutant and no significant changes in the *nem1*Δ and lipin *ned1*Δ mutants (Fig. 2A). As expected, lipin pathway deficiency resulted in a significant decrease in cellular DG and triacylglycerol (TG) levels (Fig. 2A, B). To investigate these changes at the subcellular level, we analysed the distribution of our sensors in cells lacking the Spo7-Nem1 phosphatase activity. Despite the overall similar or lower cellular abundance, PA was enriched at the INM of both *spo7*Δ and *nem1*Δ mutant cells growth in the rich YES medium (Fig. 2C, D). The DG levels at the INM were correspondingly lower (Fig. 2E, F).

**Figure 2.**
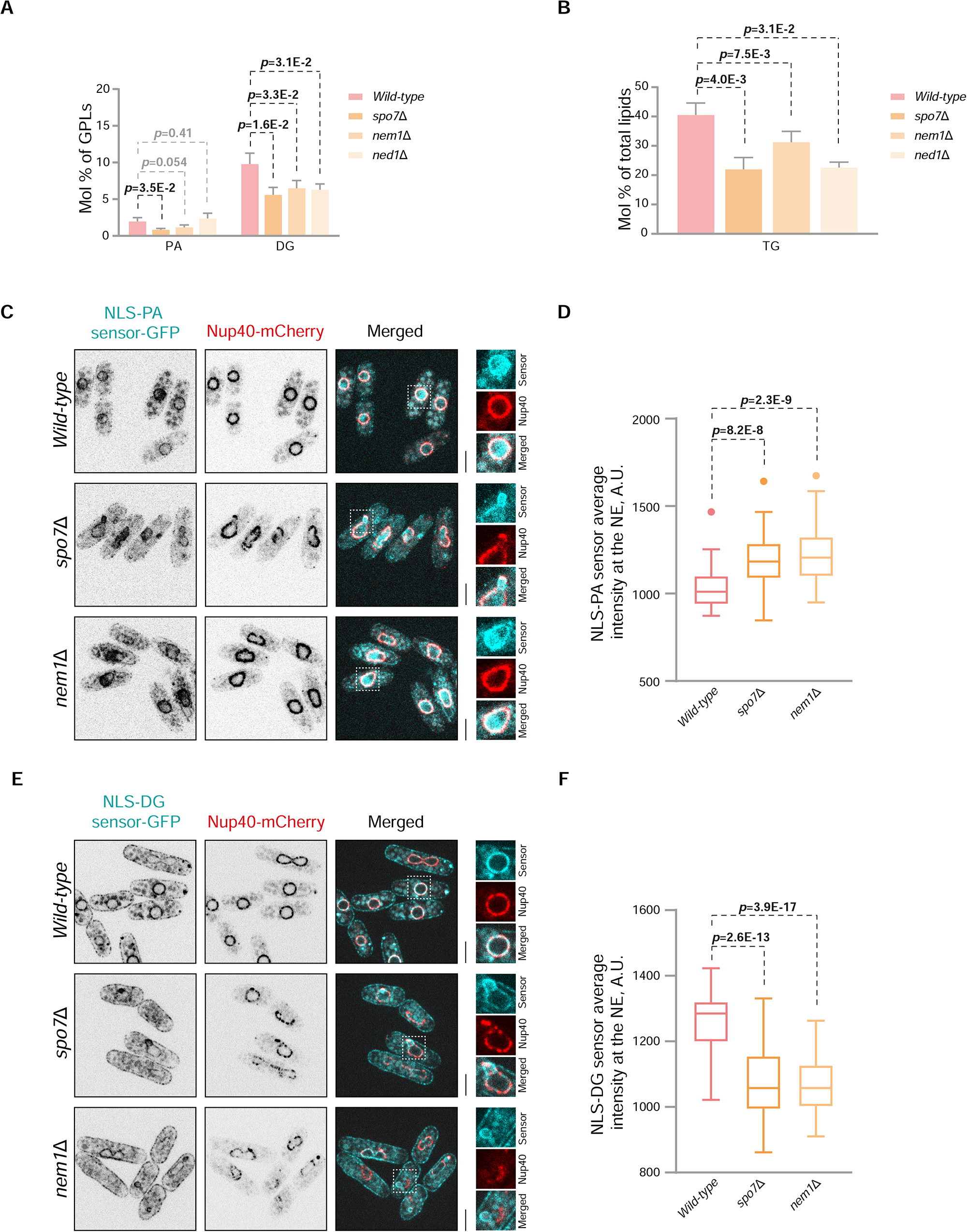
Lipid biosensors are useful to detect changes in PA and DG distribution at the subcellular level. Relative abundance of PA and DG (**A**) and TG (**B**) in *S. pombe* lipin pathway mutants. PA levels at the INM are mildly increased (**C**, **D**) and DG levels are decreased (**E**, **F**) when lipin is inactive in the absence of its activator phosphatase Spo7-Nem1. Note that whereas the NLS-DG-GFP sensor enrichment at the INM is decreased overall, we detect bright patches at the NE of some nuclei. (**C**, **E**) Single-plane spinning-disk confocal images of *S. pombe* cells expressing indicated sensors. Nucleoporin Nup40-mCherry marks the nuclear boundary. Insets represent magnified areas indicated by the dotted lines; scale bars represent 5µm. (**D**, **F**) n=50 cells, p-values are derived from the unpaired t-test.

Of note, the NLS-DG sensor was occasionally enriched at the pronounced NE ‘flares’ formed in *S. pombe spo7*Δ and *nem1*Δ mutants (Fig. 2E, insets). The DG kinase Dgk1 was not excluded from the NLS-DG sensor-rich domains (Fig. S2A, insets). These patches were found throughout the nuclear periphery, i.e., they were not associated with the nucleolus (Fig. S2B) (Witkin et al., 2012). These domains could be also induced by the major reduction in cellular DG through the overexpression of Dgk1 (Fig. S2C), or by increasing the absolute amount of NLS-DG sensor by expressing it under the strong *tdh1* promoter (Fig. S2D). We reason that such local enrichments of NLS-DG sensor may result from the imbalance between the sensor and its target lipid, and care should be taken when designing the experiments aimed at evaluating the suborganellar distribution of DG, particularly in the mutant backgrounds affecting its abundance.

Taken together, our results indicate that the newly developed genetically encoded sensors are useful as reporters to probe changes in PA and DG distribution in live fission yeast cells.

### DG at the inner nuclear membrane is depleted during closed mitosis of *S. pombe*

Time-lapse imaging showed that the fluorescence intensity of the NLS-PA sensor at the INM decreased mildly during mitosis (Fig. 3A-C). This suggested that lipin inactivation did not cause a spike in PA abundance at the NE as previously proposed for budding yeast (Han et al., 2008). Presumably, upon lipin inactivation all excess PA was diverted for GPL production, necessary for NE expansion (Makarova et al., 2016). The mild decrease could reflect the dilution effect due to the expansion in the surface area of the NE (for review see (Zhang and Oliferenko, 2013)).

**Figure 3.**
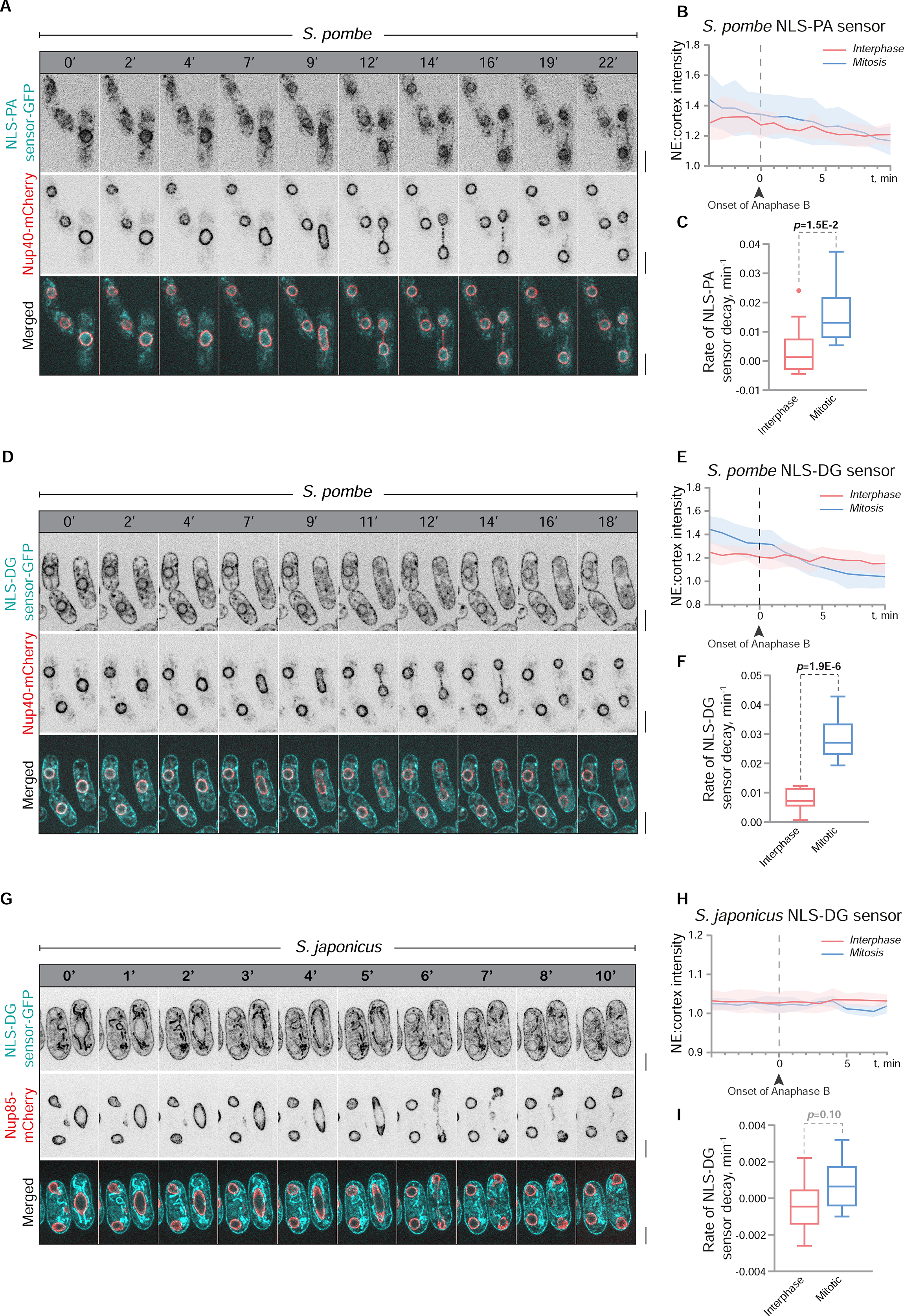
The decay of inner nuclear membrane DG during closed mitosis in *S. pombe* does not result in higher PA levels at the NE. (**A**) Time-lapse microscopy montage of mitotic *S. pombe* expressing NLS-PA sensor. (**B**) Average PA levels at the INM of *S. pombe* are mildly decreasing over the course of mitosis as compared to interphase cells. (**C**) Linear regression shows the rate of the NLS-PA sensor decay in mitotic cells is significantly different as compared to interphase cells. (**D**) Time-lapse microscopy montage of mitotic *S. pombe* expressing the NLS-DG sensor. (**E**) The NLS-DG sensor intensity at the INM rapidly decays as mitosis progresses. (**F**) The rate of decrease of the NLS-DG sensor at the INM of mitotic *S. pombe* is significantly higher than in interphase. (**G**) Time-lapse microscopy montage of the related fission yeast *S. japonicus* that does not expand the NE during mitosis. The NE breakage occurs between 5’ and 6’. (**H**) DG levels at the INM remain relatively constant throughout the semi-open mitosis of *S. japonicus*, quantified in (**I**). (**A**, **D**, **G**) Time-lapse spinning disk microscopy; shown are the maximum projections of three Z-slices; scale bars represent 5µm. Nucleoporins Nup40-mCherry (**A**, **D**) and Nup85-mCherry (**G**) mark the nuclear boundary. (**B**, **C**, **E**, **F**, **H**, **I**) n=10 cells, p-values are derived from the unpaired t-test.

Strikingly, quantitation of the NLS-DG sensor intensity over time revealed that DG levels decreased approximately at twice the rate of PA during *S. pombe* mitosis (Fig. 3D-F). Such a depletion in DG could be due, at least in part, to CDK1-dependent lipin inactivation. In fact, DG levels remained constant at the INM of mitotic *S. japonicus* (Fig. 3G-I), as lipin regulation is not tied to the cell cycle in this organism (Makarova et al., 2016). Interestingly, the subcellular enrichment of both PA and DG in *S. japonicus* was different from *S. pombe*. PA was only very weakly enriched on the NE and the cortex of *S. japonicus* (Fig. S3A), whereas DG was detected at the cortex and both sides of the NE (Fig. 3G and Fig. S3B). DG levels at the ONM of *S. japonicus* remained stable during mitosis (Fig. S3B-D), similar to its behaviour at the INM (Fig. 3G-I).

Given that *S. japonicus* does not expand the NE during mitosis, our results suggest that the depletion of DG at the INM of *S. pombe* reflects rerouting of the lipid biosynthetic flow towards GPL production necessary for nuclear membrane expansion.

### DG-to-PA conversion by Dgk1 contributes to NE expansion during ‘closed’ mitosis

We next wondered if the DG kinase Dgk1 contributed to the decrease in DG levels at the INM during mitosis of *S. pombe*. Dgk1 was previously identified in *S. cerevisiae* (Han et al., 2008), and was recently shown to display a mild under-expanded ER phenotype (Papagiannidis et al., 2021), hinting at its role in controlling ER growth. The *S. pombe* ortholog is encoded by the *ptp4/dgk1* gene (SPBC3D6.05) (referred to as Dgk1). However, the role of Dgk1 during mitosis and its effect on the subcellular distribution of PA and DG in any system remained unknown.

Consistent with previous observations in *S. cerevisiae* (Han et al., 2008), the loss of Dgk1 in *S. pombe* was able to rescue the NE and ER expansion phenotypes of the lipin pathway mutants (Fig. S4A, B), indicating the overall circuitry conservation. The loss of Dgk1 did not rescue the large lipid droplet phenotype in the lipin-deficient mutants (Fig. S4C) but restored the number of LDs to the wild-type levels (Fig. S4D).

While analysing *dgk1*Δ *S. pombe* strains we consistently noticed the presence of larger diploids in exponentially growing cultures of haploid cells. Indeed, flow cytometric analysis showed progressive accumulation of diploids in *S. pombe dgk1*Δ cultures (Fig. 4A). In *dgk1*Δ *S. japonicus*, diploidization was not detected (Fig. 4B). We reasoned that if the loss of Dgk1 resulted in a deficiency in NE expansion, the elongating spindle would experience compressive stress along its longitudinal direction, buckle and break, similarly to cells unable to produce fatty acids (Yam et al., 2011) or deficient in mitotic lipin inactivation (Makarova et al., 2016). The spindle collapse would result in the failure to divide the nucleus and diploidization. Time-lapse sequences of mitotic wild-type *S. pombe* co-expressing α-tubulin mCherry-Atb2 and the NLS-DG-GFP sensor, growing in rich medium, showed that the elongating anaphase spindles remained largely straight and DG levels at the INM decayed (Fig. 4C). Interestingly, the *dgk1*Δ *S. pombe* cells under the same conditions exhibited a range of mitotic phenotypes. Severe buckling of the mitotic spindle was observed in 31% of cells, resulting in bow-shaped nuclear intermediates during anaphase (Fig. 4D). It took longer for these nuclei to divide, and they often formed the daughter nuclei of unequal sizes (Fig. S4E). In 11% of *dgk1*Δ *S. pombe* cells, the NE failed to expand completely, and the mitotic spindle buckled and broke within the confines of the NE (Fig. 4E). Such cells may end up containing the nuclei with twice the genetic content and therefore are the likely source of a diploid cell population in *dgk1*Δ cultures. The remaining 57% of *dgk1*Δ *S. pombe* had no obvious mitotic phenotypes and divided normally (Fig. S4F, G).

**Figure 4.**
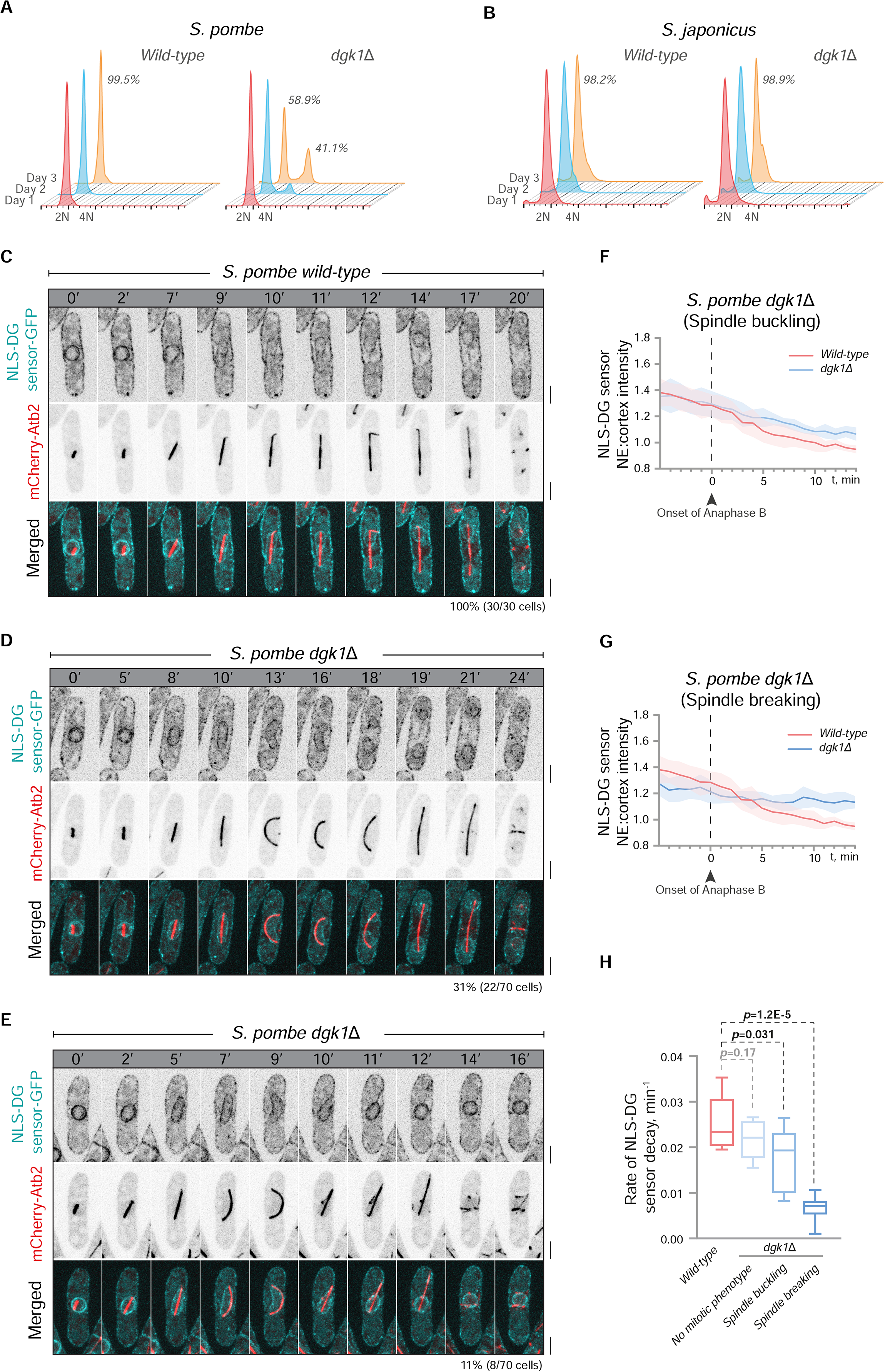
The DG kinase Dgk1 is essential for maintaining mitotic fidelity in *S. pombe*. (**A**) Diploids accumulate upon prolonged culturing of *dgk1*Δ *S. pombe*, as shown by flow cytometric analysis. (**B**) In *S. japonicus*, the loss of Dgk1 does not result in diploidization. (**C**-**E**) Representative time-lapse microscopy montages show a range of anaphase phenotypes in *dgk1*Δ *S. pombe*. (**C**) In the wild-type, the spindle remains relatively straight throughout mitosis, indicating that it is not constrained by the NE. (**D**) 31% of *dgk1*Δ *S. pombe* exhibit anaphase spindle buckling with eventual relaxation, whereas (**E**) 11% break spindles following their compression within the confines of the NE. The decrease in the NLS-DG sensor intensity in cells exhibiting spindle buckling (**F**) and breaking (**G**) phenotypes is compared against the wild-type. (**H**) The rate of decrease of the NLS-DG sensor intensity at the INM is predictive of the severity of mitotic phenotype in *dgk1*Δ *S. pombe*. (**A**-**B**) Shown are representative graphs from at least three biological repeats. (**C**-**E**) Time-lapse spinning disk microscopy; shown are the maximum projections of three Z-slices; scale bars represent 5µm. α-tubulin mCherry-Atb2 marks the mitotic spindle. (**F**-**H**) n=8 cells, p-values are derived from the unpaired t-test.

Strikingly, we observed a correlation between the rates of DG decay at the INM and the severity of the mitotic phenotypes in *dgk1*Δ *S. pombe* cells (Fig. 4F-H and Fig. S4G). Whereas DG levels were depleted similarly to the wild-type in the normally dividing *dgk1*Δ mutants (Fig. S4G), they decayed less in cells with buckling spindles (Fig. 4F) and remained constant in those cells that failed to expand the NE (Fig. 4G).

Thus, both the activity of Dgk1 as well as concurrent inactivation of lipin contribute to the decrease in DG levels at the NE during closed mitosis in *S. pombe*. Importantly, this indicates that Dgk1 activity promotes NE expansion required for ‘closed’ mitosis. The fact that DG may decay at different rates in mitotic *dgk1*Δ cells yielding distinct nuclear division phenotypes suggests an underlying metabolic heterogeneity even within isogenic cellular populations.

### The balance between DG and PA but not the elevated levels of PA is the hallmark of NE expansion in *S. pombe*

Given that the lack of Dgk1 counteracted excessive NE-ER expansion in the lipin pathway mutants (Fig. S4A, B), we wondered if the loss of lipin activity resulting in a drop in DG and an increase in PA at the NE (Fig. 2D, F), may in turn alleviate insufficient mitotic NE expansion observed in *dgk1*Δ mutant cells. Intriguingly, the *spo7*Δ *dgk1*Δ double mutants displayed normal mitosis and a decay of DG levels at the NE similar to the wild-type (Fig. 5A, B, see also Fig. S5A, B for *nem1*Δ *dgk1*Δ data). Moreover, we have observed a significant reduction in the diploidization in the cultures of the double mutants compared to the *dgk1*Δ strain (Fig. 5C and Fig. S5C).

**Figure 5.**
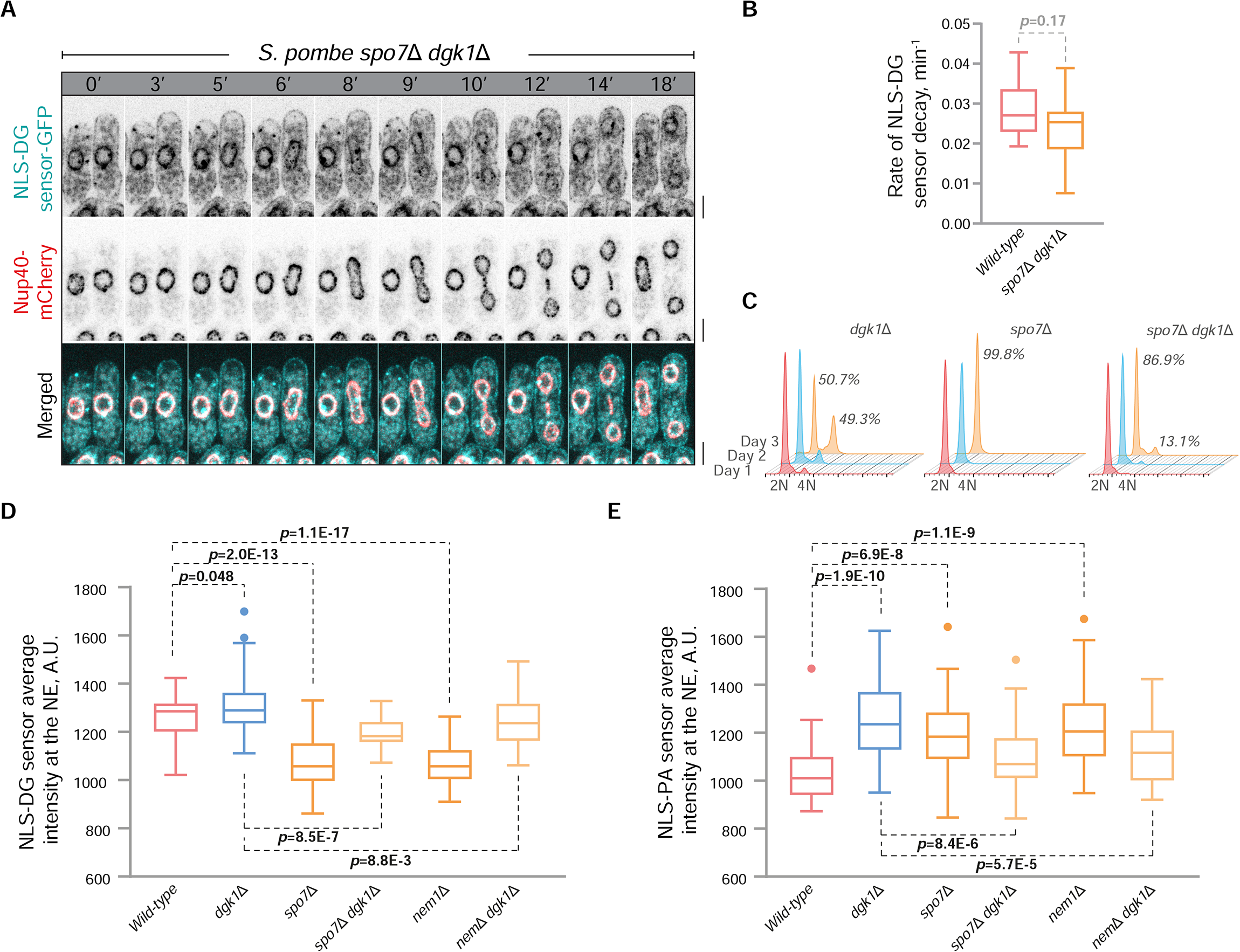
Changes in the PA-to-DG ratio at the INM are consistent with NE expansion in *S. pombe.* (A) Time-lapse microscopy of the *spo7*Δ *dgk1*Δ double mutant undergoing mitosis shows a decay of the NLS-DG sensor intensity at the INM, similar to the wild-type. Shown are the maximum projections of three Z-slices. Nucleoporin Nup40-mCherry marks the nuclear boundary; scale bars represent 5µm. Quantification of the rate of decay is shown in (**B**). (**C**) Flow cytometry shows that the *spo7*Δ *dgk1*Δ double mutant has a lower proportion of diploids in the population as compared to the *dgk1*Δ cells. Quantification of the levels of DG (**D**) and PA (**E**) at the INM in the indicated genotypes of interphase *S. pombe*. The lack of Dgk1 results in an increase in PA and a mild increase in DG at the INM, which can be reduced by the deletion of the activator of lipin, the phosphatase Spo7-Nem1. (**B**) n=10 cells; (**D**, **E**) n=50 cells; (**B**, **D**, **E**) p-values are derived from the unpaired t-test.

As expected, Dgk1 deficiency resulted in a mild increase in DG levels at the NE (Fig. 5D). Perhaps less intuitively, the PA levels increased substantially in this genetic background, similarly to the lipin pathway mutants (Fig. 5E). This argues that increasing PA levels alone is not sufficient for NE expansion (Han et al., 2008). Instead, we observe changes in the relative abundance of PA and DG at the NE. Indeed, PA increase in the lipin pathway mutants is accompanied by the drop in DG, whereas this is not the case in cells lacking Dgk1. The double mutants lacking both lipin and Dgk1 activities largely restore the relative abundances of PA and DG (Fig. 5D, E) and show normal NE morphology (Fig. S4A, B).

Taken together, our results suggest that regulation of DG levels at the INM is important in controlling NE expansion. Yet, how is DG at the NE removed in cells that lack Dgk1 but do not display any mitotic phenotype?

### The Kennedy pathway contributes strongly to glycerophospholipid synthesis required for NE expansion during ‘closed’ mitosis

In addition to Dgk1-dependent rerouting of DG towards PA for *de novo* glycerophospholipid synthesis, DG might be also utilized directly by the precursor-dependent Kennedy pathway to produce glycerophospholipids (Carman and Han, 2009b). Our observations suggest that the contribution of the Kennedy pathway is critical for proper nuclear membrane expansion during mitosis. First, *dgk1*Δ mutant cultures grown in the chemically defined medium (EMM) that does not contain choline and ethanolamine (Cho/Etn) precursors required for the Kennedy pathway exhibited a high incidence of cells with unequally dividing nuclei and complete failure in nuclear division (‘cut’ phenotype) (Fig. 6A-C). Many cells in these cultures became non-viable (Fig. 6D). Second, whereas PA was highly abundant at the INM of *dgk1*Δ cells grown in the presence of the Kennedy pathway (the rich YES medium or EMM+Cho/Etn), we did not observe any enrichment in the absence of the Kennedy pathway precursors (Fig. 6E, compare with Fig. 5E). This suggests that rerouting of DG to GPL biosynthesis via the Kennedy pathway might decrease the consumption of PA via the CDP-DG and/or lipin pathways. Differential activity of the DG-consuming Kennedy pathway within a population of cells could be an underlying cause for the heterogeneity in cellular phenotypes associated with the loss of Dgk1.

**Figure 6.**
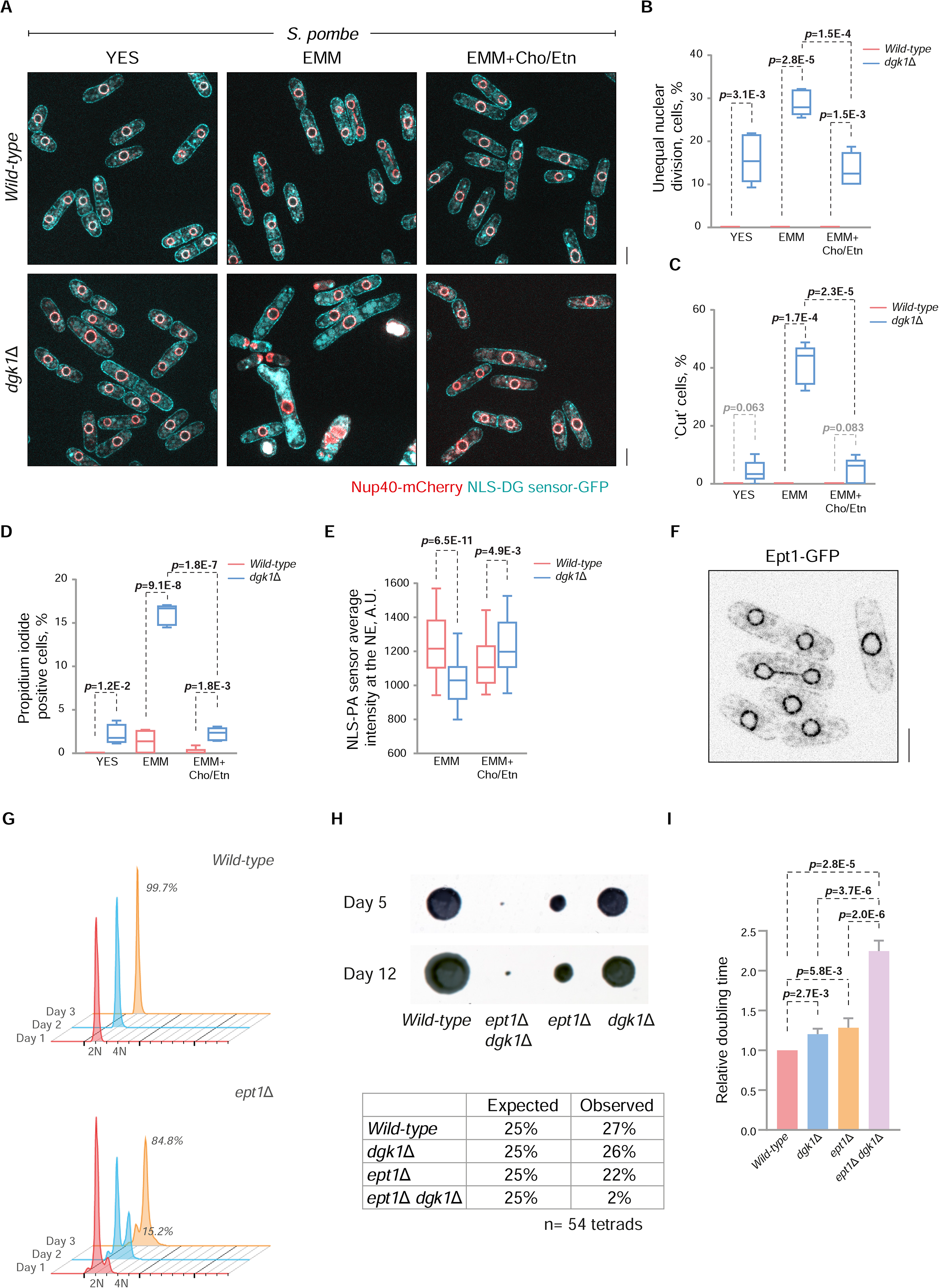
The Kennedy pathway of GPL synthesis collaborates with Dgk1 in sustaining NE expansion required for closed mitosis in *S. pombe*. (**A**) In the absence of the Kennedy pathway precursors, *dgk1*Δ cells exhibit a substantial increase in uneven nuclear divisions (quantitation in **B**) and mitotic failure (‘cut’ phenotype) (quantitation in **C**). Shown are the single-plane spinning-disk confocal images of *S. pombe* cells co-expressing NLS-DG-GFP sensor and Nup40-mCherry. Scale bars represent 5µm. (**D**) High levels of cell death was detected by propidium iodide staining in *S. pombe dgk1*Δ cells cultured in the absence of the Kennedy pathway precursors. (**E**) Quantification of the levels of PA at the INM in wild-type and *dgk1*Δ cells grown in the indicated medium. The DG-dependent Kennedy pathway reduces consumption of PA at the INM. (**F**) Single-plane spinning-disk confocal images show Ept1-GFP localization at the NE of *S. pombe*. (**G**) Diploids accumulate upon culturing of the Kennedy pathway deficient *ept1*Δ mutant strain of *S. pombe*, as shown by flow cytometric analysis. (**H**) Tetrad analysis reveals the semi-lethality of the *ept1*Δ *dgk1*Δ double mutant. Shown here is a representative tetrad grown on rich YES agar on day 5 and day 12 after dissection. (**I**) Relative growth rates of the indicated genotypes in rich YES media. (**B**, **C**) n=4 biological repeats, ≥10 septated cells each; (**D**) n=5 biological repeats, ≥50 cells each; (**E**) n=50 cells; (**H**) n=54 tetrads; (**I**) n=5 technical repeats. (**B**-**E**) p-values are derived from the unpaired t-test.

Underscoring that the *de novo* and the Kennedy pathways both contribute to GPL synthesis, we detected a decrease in NE-ER proliferation in the minimal EMM medium, as compared to either EMM containing Cho/Etn precursors or rich YES medium (Fig. S6A, B). As DG synthesis is decreased in the lipin-deficient mutants, the Kennedy pathway presumably becomes sufficient for consumption of DG at the NE, explaining the mitotic DG decay observed in *spo7*Δ *dgk1*Δ double mutants (Fig. 5A, B; see also Fig. S5A, B for *nem1*Δ *dgk1*Δ data).

Ept1 (SPAC22A12.10) is the diacylglycerol cholinephosphotransferase / ethanolaminephosphotransferase that catalyses the final reaction of the Kennedy pathway in fission yeast. Ept1-GFP is enriched at the NE of *S. pombe* throughout the cell cycle, suggesting that the DG-dependent GPL biosynthesis may occur at this location (Fig. 6F). In order to test directly if the Kennedy pathway contributes to NE expansion, we generated a mutant strain lacking Ept1. The *ept1*Δ mutants exhibited striking diploidization with a majority of the population having double the genetic content within three days of consecutive culture (Fig. 6G). The introduction of the tagged α-tubulin mCherry-Atb2 triggered a much higher incidence of mitotic failure, with mutant cultures swept by diploids even after short period of culturing (Fig. S6C). Importantly, the *ept1*Δ *dgk1*Δ double mutant phenotype was sub-lethal, as observed through genetic crosses and phase-contrast microscopy (Fig. 6H and S6D). Those rare *ept1*Δ *dgk1*Δ double mutants that we were able to recover exhibited severe delay in growth, as compared to the wild-type and the single *ept1*Δ and *dgk1*Δ mutants (Fig. 6I).

In summary, our results show that controlling the utilization of DG at the INM is important for the regulation of NE expansion in *S. pombe*. Both the Dgk1-dependent DG-to-PA conversion and the Kennedy pathway collaborate to channel DG to GPL production required for a mitotic spike in NE growth.

## Discussion

Our results show that NE expansion in *S. pombe* appears to require a pronounced drop in DG levels at the INM. This DG depletion is the result of several enzymatic reactions. Lipin that synthesizes DG from PA is inactivated at mitotic entry by CDK1 (Carman and Han, 2009a; Makarova et al., 2016), reducing DG inflow. At the same time, DG can be either phosphorylated by Dgk1 to produce PA (Han et al., 2008; Kwiatek et al., 2020) or directly used for GPL biosynthesis through the Kennedy pathway (Carman and Han, 2009b). As we do not observe PA levels spiking at the NE at mitosis, it is likely that the Dgk1-produced PA is being used up for the synthesis of membrane lipids through the *de novo* CDP-DG route (Han et al., 2008; Kwiatek et al., 2020) (Fig. 7).

**Figure 7.**
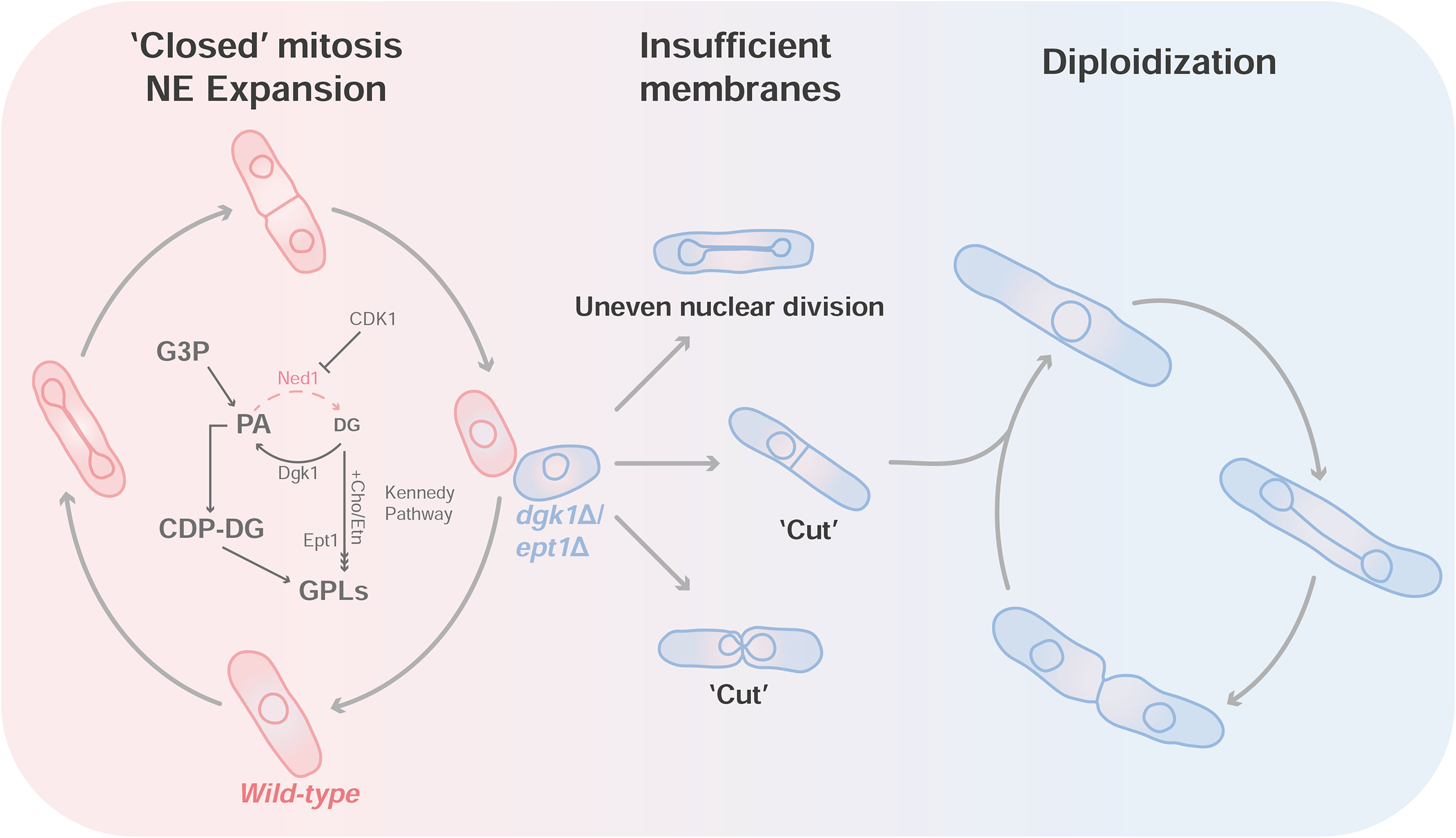
Summary diagram. A summary of our hypothesis on how PA and DG flux during NE expansion is controlled in *S. pombe.* In wild-type *S. pombe*, the inactivation of lipin Ned1 by CDK1, together with Dgk1 activity and the Kennedy pathway, channel DG towards GPL biosynthesis. In the absence of Dgk1, insufficient depletion of DG results in reduced membrane availability or/and changes in the biophysical properties of the membrane. When Dgk1 is inactive, cells rely on the Kennedy pathway-dependent DG consumption to produce GPLs. Cell-to-cell heterogeneity in the activity of the Kennedy pathway results in mitotic phenotypes such as uneven nuclear division or chromosome segregation defects (‘cut’ phenotype). Many cells end up with twice the genetic content, forming diploids.

The lipidomics analysis of the *S. cerevisiae* lipin mutant showed an increase in cellular PA levels (Han et al., 2008). Such an increase in PA, a conically shaped lipid capable of generating negative membrane curvature (Zhukovsky et al., 2019), was inferred to modify nuclear membrane properties and lead to the transcriptional upregulation of GPL biosynthetic genes through the Opi1p-Ino2p-Ino4p regulation circuitry (Carman and Henry, 2007; Han et al., 2008), although the latter is not conserved in fission yeasts (Rhind et al., 2011; Rutherford et al., 2022). Of note, PA does not seem to increase neither on global scale in the lipin-deficient *S. pombe* cells (Fig. 2A), nor at the NE during mitosis, when lipin function is inhibited by CDK1 (Fig. 3A, B and (Makarova et al., 2016)). It also does not increase in the budding yeast *spo7*Δ and *nem1*Δ mutants despite the block to lipin activity (Papagiannidis et al., 2021).

Similar to PA, DG has negative spontaneous curvature and was shown to promote fusion of biological membranes, NE assembly, and LD formation (Dumas et al., 2010; Domart et al., 2012; Miner et al., 2017; Choudhary et al., 2018; Chung et al., 2018). It is possible that in addition to fuelling membrane lipid synthesis, depletion of DG or changes in the PA-to-DG ratio may modify the biophysical properties of the NE priming it for expansion.

However, we favour the possibility that rather than having functional implications for NE remodelling, the PA-to-DG ratio at the NE simply reflects the utilization of these lipids by different biosynthetic pathways. Both lipids are precursors for the biosynthesis of other lipid species. GPLs are mainly synthesized through two pathways, the *de novo* CDP-DG-dependent route, which uses PA, and the precursor-dependent Kennedy pathway, which uses DG (McMaster and Bell, 1994; Gibellini and Smith, 2010). In addition, DG can be used for the production of the storage lipid TG (Carman and Henry, 2007; Carman and Han, 2009b; Holic et al., 2020). The differential contribution of these biosynthetic pathways may manifest as changes in the steady state PA-to-DG ratio.

The range of mitotic phenotypes within isogenic populations of Dgk1-deficient cells grown in rich media suggests an underlying metabolic heterogeneity. This may arise due to a number of reasons, e.g., differences in the Kennedy pathway precursor uptake (Gibellini and Smith, 2010), the differential activities of biosynthetic pathways (Stewart-Ornstein et al., 2012), or stochasticity in gene expression (Kaern et al., 2005; Raj and van Oudenaarden, 2008). The phenotypic heterogeneity largely collapses when the Kennedy pathway is disabled either by withdrawal of precursors or the loss of Ept1 (Fig. 6), suggesting that the Kennedy pathway activity may indeed show cell-to-cell variability.

Both Dgk1 and the Kennedy pathway have a role in NE expansion that enables ‘closed’ mitosis and, hence, the maintenance of mitotic fidelity in *S. pombe* (Fig. 4-6; see a diagram in Fig. 7). Interestingly, a functional genomics screen has linked the loss of Dgk1 to meiotic chromosome segregation defects (Blyth et al., 2018). These phenotypes were speculated to be a result of global deregulation in lipid synthesis (Holic et al., 2020). Our data suggest that one possible alternative explanation could be membrane limitation during meiotic remodelling of the NE.

The bulk of the NE membrane expansion during closed mitosis likely occurs *in situ* or at the ER domain close to the NE (Fig. 6F and Fig. S2A). Our results may have implications for mitosis in other organisms. A range of mitotic strategies in nature spans from completely ‘closed’ to completely ‘open’ mitosis, when the NE breaks down in prophase and reforms around the segregated chromosomes upon mitotic exit (Makarova and Oliferenko, 2016). Recent evidence points out at the importance of regulating GPL synthesis for NE reformation and other dynamic events at the NE in metazoans (Bahmanyar and Schlieker, 2020), and it would be of interest to address if the Kennedy pathway or the control of DG-to-PA conversion by diacylglycerol kinases are also critical for these processes.

Advancements in the system-level analyses of lipids have allowed greater scrutiny of their roles in various cellular processes (Han, 2016; Kofeler et al., 2021). Yet, these approaches struggle with informing on lipid dynamics with fine spatiotemporal resolution. Complementing lipidomics and genetics approaches with a set of newly developed lipid sensors for *S. pombe* have allowed us to track changes in the spatiotemporal distribution of PA and DG with subcellular resolution and formulate and test a set of hypotheses on the roles of PA to DG interconversion during mitotic NE remodelling. We believe that these sensors will become an effective and widely used tool for the study of dynamic membrane processes in fission yeasts.

## Materials and methods

### Strains, media, and molecular biology methods

*S. pombe* and *S. japonicus* strains used in this study are listed in Table 1. Standard fission yeast media and methods were used (Moreno et al., 1991; Aoki et al., 2010). All experiments were performed using the rich non-defined YES medium with the exception of lipidomics, thiamine-repressible *nmt1* (Maundrell, 1990) overexpression of Dgk1, and the Kennedy pathway-related experiments shown in Fig. 6F-J and Fig. S6D, E, which were performed using chemically-defined Edinburgh Minimal Medium (EMM). All strains were grown at 30°C unless otherwise specified, in temperature-controlled 200rpm shaking incubators. All mating was performed on SPA medium and spores dissected on YES agar plates. Homozygous *S. pombe* diploids are generated by *ade6-M210*/*ade6-M216* heteroallelic complementation (Ekwall and Thon, 2017). Choline and ethanolamine (Sigma-Aldrich) were added to EMM at final concentrations of 1mM.

| Strain | Source | Identifier | Figures |
| --- | --- | --- | --- |
| *ura4+::cdc15<sup>S.p.</sup> promoter*-PA sensor (Opi1<sup>111-189, G120W</sup> 2xAH)-GST-GFP *ade6-704 ura4-294 leu1-32 h-* | This study | SO8772 | 1B, S1B |
| *ura4+::cdc15<sup>S.p.</sup> promoter*-NLS PA sensor (SV40 NLS-Opi1<sup>111-189, G120W</sup> 2xAH-SV40 NLS)-GST-GFP *ade6-704 ura4-294 leu1-32 h-* | This study | SO8760 | 1B, S1B |
| *ura4+::cdc15<sup>S.p.</sup> promoter*-PA sensor binding mutant (Opi1<sup>111-189, G120W, L124R, Y127A, L129R, M131A, I133R, K136A, K137A, R138A</sup> 2xAH)-GST-GFP *ade6-704 ura4-294 leu1-32 h-* | This study | SO8815 | 1C |
| *ura4+::cdc15<sup>S.p.</sup> promoter*-NLS PA sensor binding mutant (SV40 NLS-Opi1<sup>111-189, G120W, L124R, Y127A, L129R, M131A, I133R, K136A, K137A, R138A</sup> 2xAH-SV40 NLS)-GST-GFP *ade6-704 ura4-294 leu1-32 h-* | This study | SO8817 | 1C |
| *ura4+::cdc15<sup>S.p.</sup> promoter*-DG sensor (2x PKCβ DG binding domain)-GST-GFP *ade6-704 ura4-294 leu1-32 h-* | This study | SO8766 | 1D, S1D |
| *ura4+::cdc15<sup>S.p.</sup> promoter*-NLS DG sensor (SV40 NLS-2x PKCβ DG binding domain-SV40 NLS)-GST-GFP *ade6-704 ura4-294 leu1-32 h-* | This study | SO8767 | 1D, S1D, S2D |
| *ura4+::cdc15<sup>S.p.</sup> promoter*-DG sensor binding mutant (2x PKCβ DG binding domain<sup>Q63E, Q128E</sup>)-GST-GFP *ade6-704 ura4-294 leu1-32 h-* | This study | SO8819 | 1E |
| *ura4+::cdc15<sup>S.p.</sup> promoter*- NLS DG sensor binding mutant (SV40 NLS-2x PKCβ DG binding domain<sup>Q63E, Q128E</sup>-SV40 NLS)-GST-GFP *ade6-704 ura4-294 leu1-32 h-* | This study | SO8887 | 1E |
| *leu1+::bip1<sup>S.p.</sup> promoter*-mCherry-ADEL *ura4+::cdc15<sup>S.p.</sup> promoter*-DG sensor (2x PKCβ DG binding domain)-GST-GFP *ade6-704 ura4-294 leu1-32 h-* | This study | SO8854 | 1F |
| *scs2Δ::ura4+ scs22Δ::ura4+ leu1+::bip1<sup>S.p.</sup> promoter*-mCherry-ADEL *ura4+::cdc15<sup>S.p.</sup> promoter*-DG sensor (2x PKCβ DG binding domain)-GST-GFP | This study | SO8844 | 1F |
| *leu1+::pREP1-Dgk1 ura4+::cdc15<sup>S.p.</sup> promoter*-PA sensor (Opi1<sup>111-189, G120W</sup> 2xAH)-GST-GFP *ade6-704 ura4-294 leu1-32 h-* | This study | SO9014 | 1G, 1H |
| *leu1+::pREP1-Dgk1<sup>D119A</sup> ura4+::cdc15<sup>S.p.</sup> promoter*-PA sensor (Opi1<sup>111-189, G120W</sup> 2xAH)-GST-GFP *ade6-704 ura4-294 leu1-32 h-* | This study | SO9018 | 1G, 1H |
| *leu1+::pREP1-Dgk1 ura4+::cdc15<sup>S.p.</sup> promoter*-DG sensor (2x PKCβ DG binding domain)-GST-GFP *ade6-704 ura4-294 leu1-32 h-* | This study | SO9016 | 1G, 1I |
| *leu1+::pREP1-Dgk1<sup>D119A</sup> ura4+::cdc15<sup>S.p.</sup> promoter*-DG sensor (2x PKCβ DG binding domain)-GST-GFP *ade6-704 ura4-294 leu1-32 h-* | This study | SO9020 | 1G, 1I |
| *ura4+::cdc15<sup>S.p.</sup> promoter*-PA sensor (Opi1<sup>103-189</sup> 1xAH)-GST-GFP *ade6-704 ura4-294 leu1-32 h-* | This study | SO9097 | S1B |
| *ura4+::cdc15<sup>S.p.</sup> promoter*-PA sensor (Opi1<sup>111-189</sup> 2xAH)-GST-GFP *ade6-704 ura4-294 leu1-32 h-* | This study | SO8777 | S1B |
| *ura4+::cdc15<sup>S.p.</sup> promoter*-PA sensor (Opi1<sup>111-189, G120W</sup> 1xAH)-GST-GFP *ade6-704 ura4-294 leu1-32 h-* | This study | SO8784 | S1B |
| *ura4+::cdc15<sup>S.p.</sup> promoter*-NLS PA sensor (SV40 NLS-Opi1<sup>103-189</sup> 1xAH-SV40 NLS)-GST-GFP *ade6-704 ura4-294 leu1-32 h-* | This study | SO9098 | S1B |
| *ura4+::cdc15<sup>S.p.</sup> promoter*-NLS PA sensor (SV40 NLS-Opi1<sup>111-189</sup> 2xAH-SV40 NLS)-GST-GFP *ade6-704 ura4-294 leu1-32 h-* | This study | SO8778 | S1B |
| *ura4+::cdc15<sup>S.p.</sup> promoter*-NLS PA sensor (SV40 NLS-Opi1<sup>111-189, G120W</sup> 1xAH-SV40 NLS)-GST-GFP *ade6-704 ura4-294 leu1-32 h-* | This study | SO8785 | S1B |
| *ura4+::cdc15<sup>S.p.</sup> promoter*-DG sensor (1x PKCβ DG binding domain)-GST-GFP *ade6-704 ura4-294 leu1-32 h-* | This study | SO9099 | S1D |
| *ura4+::cdc15<sup>S.p.</sup> promoter*- NLS DG sensor (SV40 NLS-1x PKCβ DG binding domain-SV40 NLS)-GST-GFP *ade6-704 ura4-294 leu1-32 h-* | This study | SO9100 | S1D |
| Wild-type 972 *h-* | Lab collection | SO8087 | 2A, 2B |
| *spo7Δ::ura4+* | Lab collection | SO8109 | 2A, 2B, 5C |
| *nem1Δ::kanR h-* | This study | SO8122 | 2A, 2B, S5C |
| *ned1Δ::ura4+* | Lab collection | SO8110 | 2A, 2B |
| *nup40-mCherry::ura4+ ura4+::cdc15<sup>S.p.</sup> promoter*-NLS PA sensor (SV40 NLS-Opi1<sup>111-189, G120W</sup> 2xAH-SV40 NLS)-GST-GFP | This study | SO8872 | 2C, 2D, 3A-C, 5E, 6E |
| *spo7Δ::kanR nup40-mCherry::ura4+ ura4+::cdc15<sup>S.p.</sup> promoter*-NLS PA sensor (SV40 NLS-Opi1<sup>111-189, G120W</sup> 2xAH-SV40 NLS)-GST-GFP | This study | SO8888 | 2C, 2D, 5E |
| *nem1Δ::kanR nup40-mCherry::ura4+ ura4+::cdc15<sup>S.p.</sup> promoter*-NLS PA sensor (SV40 NLS-Opi1<sup>111-189, G120W</sup> 2xAH-SV40 NLS)-GST-GFP | This study | SO8889 | 2C, 2D, 5E |
| *nup40-mCherry::ura4+ ura4+::cdc15<sup>S.p.</sup> promoter*-NLS DG sensor (SV40 NLS-2x PKCβ DG binding domain-SV40 NLS)-GST-GFP *h+* | This study | SO8874 | 2E, 2F, 3D-F, 5B, 5D, S5B, 6A-C |
| *spo7Δ::kanR nup40-mCherry::ura4+ ura4+::cdc15<sup>S.p.</sup> promoter*-NLS DG sensor (SV40 NLS-2x PKCβ DG binding domain-SV40 NLS)-GST-GFP *h+* | This study | SO8890 | 2E, 2F, 5D |
| *nem1Δ::kanR nup40-mCherry::ura4+ ura4+::cdc15<sup>S.p.</sup> promoter*-NLS DG sensor (SV40 NLS-2x PKCβ DG binding domain-SV40 NLS)-GST-GFP *h-* | This study | SO8891 | 2E, 2F, 5D |
| *dgk1-GFP::natR ura4+::cdc15<sup>S.p.</sup> promoter-NLS DG sensor (SV40 NLS-2x PKCβ DG binding domain-SV40 NLS)-GST-GFP* | This study | SO9188 | S2A |
| *spo7Δ::kanR dgk1-GFP::natR ura4+::cdc15<sup>S.p.</sup> promoter-NLS DG sensor (SV40 NLS-2x PKCβ DG binding domain-SV40 NLS)-GST-mCherry* | This study | SO9207 | S2A |
| *nem1Δ::kanR dgk1-GFP::natR ura4+::cdc15<sup>S.p.</sup> promoter-NLS DG sensor (SV40 NLS-2x PKCβ DG binding domain-SV40 NLS)-GST-mCherry* | This study | SO9208 | S2A |
| *bop1-mCherry::ura4+ ura4+::cdc15<sup>S.p.</sup> promoter-NLS DG sensor (SV40 NLS-2x PKCβ DG binding domain-SV40 NLS)-GST-GFP* | This study | SO8919 | S2B |
| *spo7Δ::kanR bop1-mCherry::ura4+ ura4+::cdc15<sup>S.p.</sup> promoter-NLS DG sensor (SV40 NLS-2x PKCβ DG binding domain-SV40 NLS) NLS-GST-GFP* | This study | SO8917 | S2B |
| *nem1Δ::kanR bop1-mCherry::ura4+ ura4+::cdc15<sup>S.p.</sup> promoter-NLS DG sensor (SV40 NLS-2x PKCβ DG binding domain-SV40 NLS)-GST-GFP* | This study | SO9093 | S2B |
| *leu1+::pREP1-Dgk1 nup40-mCherry::ura4+ ura4+::cdc15<sup>S.p.</sup> promoter-NLS DG sensor (SV40 NLS-2x PKCβ DG binding domain-SV40 NLS)-GST-GFP h-* | This study | SO9017 | S2C |
| *leu1+::pREP1-Dgk1<sup>D119A</sup> nup40-mCherry::ura4+ ura4+::cdc15<sup>S.p.</sup> promoter-NLS DG sensor (SV40 NLS-2x PKCβ DG binding domain-SV40 NLS)-GST-GFP h-* | This study | SO9021 | S2C |
| *ura4+::tdh1<sup>S.p.</sup> promoter-NLS DG sensor (SV40 NLS-2x PKCβ DG binding domain-SV40 NLS)-GST-GFP ade6-704 ura4-294 leu1-32 h-* | This study | SO9086 | S2D |
| *nup85-mCherry::ura4+ ura4+::cdc15<sup>S.p.</sup> promoter-NLS DG sensor (SV40 NLS-2x PKCβ DG binding domain-SV40 NLS)-GFP h+* | This study | SOJ5220 | 3G-I |
| *nup85-mCherry::ura4+ ura4+::cdc15<sup>S.p.</sup> promoter-PA sensor (Opi1<sup>111-189, G120W</sup> 2xAH)-GFP h+* | This study | SOJ5218 | S3A |
| *nup85-mCherry::ura4+ ura4+::cdc15<sup>S.p.</sup> promoter-NLS PA sensor (SV40 NLS-Opi1<sup>111-189, G120W</sup> 2xAH-SV40 NLS)-GFP h+* | This study | SOJ5219 | S3A |
| *nup85-mCherry::ura4+ ura4+::cdc15<sup>S.p.</sup> promoter-DG sensor (2x PKCβ DG binding domain)-GFP* | This study | SOJ5238 | S3B-D |
| Wild-type *ade6-210 ura4-D18 leu1-32 h+* | Lab collection | SO2865 | 4A, 6D, 6G, 6I, S6C |
| *dgk1Δ::natR ade6-216 ura4-D18 leu1-32 h-* | This study | SO8928 | 4A, 5C, S5C, 6D, 6I |
| Wild-type *ura4sj-D3 ade6sj-domE matsj-P2028 h-* | Lab collection | SOJ11 | 4B |
| *dgk1Δ::natR ura4sj-D3 ade6sj-domE matsj-P2028 h-* | This study | SOJ4942 | 4B |
| *mCherry-atb2::hphR ura4+::cdc15<sup>S.p.</sup> promoter-NLS DG sensor (SV40 NLS-2x PKCβ DG binding domain-SV40 NLS)-GST-GFP ade6-216 h-* | This study | SO9058 | 4C, 4F-H, S4G, S6C |
| *dgk1Δ::natR mCherry-atb2::hphR ura4+::cdc15<sup>S.p.</sup> promoter-NLS DG sensor (SV40 NLS-2x PKCβ DG binding domain-SV40 NLS)-GST-GFP ade6-216 h-* | This study | SO9059 | 4D-H, S4E-G |
| *leu1+::bip1<sup>S.p.</sup> promoter-mCherry-ADEL h+* | Lab collection | SO5609 | S4A-D, S6A, S6B |
| *dgk1Δ::natR leu1+::bip1<sup>S.p.</sup> promoter-mCherry-ADEL h-* | This study | SO8930 | S4A-D |
| *spo7Δ::ura4+ leu1+::bip1<sup>S.p.</sup> promoter-mCherry-ADEL* | This study | SO8733 | S4A-D, S6A, S6B |
| *dgk1Δ::leu1+ spo7Δ::ura4+ leu1+::bip1<sup>S.p.</sup> promoter-mCherry-ADEL* | This study | SO8902 | S4A-D |
| *nem1Δ::kanR leu1+::bip1<sup>S.p.</sup> promoter-mCherry-ADEL* | This study | SO8734 | S4A-D, S6A, S6B |
| *dgk1Δ::leu1+ nem1Δ::kanR leu1+::bip1<sup>S.p.</sup> promoter-mCherry-ADEL* | This study | SO8903 | S4A-D |
| *spo7Δ::kanR dgk1Δ::leu1+ nup40-mCherry::ura4+ ura4+::cdc15<sup>S.p.</sup> promoter-NLS DG sensor (SV40 NLS-2x PKCβ DG binding domain-SV40 NLS)-GST-GFP* | This study | SO9094 | 5A, 5B, 5D |
| *dgk1Δ::leu1+ spo7Δ::kanR* | This study | SO9269 | 5C |
| *dgk1Δ::leu1+ nup40-mCherry::ura4+ cdc15<sup>S.p.</sup> promoter-NLS DG sensor (SV40 NLS-2x PKCβ DG binding domain-SV40 NLS)-GST-GFP* | This study | SO8915 | 5D, 6A-C |
| *nem1Δ::kanR dgk1Δ::leu1+ nup40-mCherry::ura4+ ura4+::cdc15<sup>S.p.</sup> promoter-NLS DG sensor (SV40 NLS-2x PKCβ DG binding domain-SV40 NLS)-GST-GFP* | This study | SO9095 | 5D, S5A, S5B |
| *dgk1Δ::natR nup40-mCherry::ura4+ ura4+::cdc15<sup>S.p.</sup> promoter-NLS PA sensor (SV40 NLS-Opi1<sup>111-189, G120W</sup> 2xAH-SV40 NLS)-GST-GFP* | This study | SO8995 | 5E, 6E |
| *spo7Δ::kanR dgk1Δ::leu1+ nup40-mCherry::ura4+ ura4+::cdc15<sup>S.p.</sup> promoter-NLS PA sensor (SV40 NLS-Opi1<sup>111-189, G120W</sup> 2xAH-SV40 NLS)-GST-GFP* | This study | SO9261 | 5E |
| *nem1Δ::kanR dgk1Δ::leu1+ nup40-mCherry::ura4+ ura4+::cdc15<sup>S.p.</sup> promoter-NLS PA sensor (SV40 NLS-Opi1<sup>111-189, G120W</sup> 2xAH-SV40 NLS)-GST-GFP* | This study | SO9262 | 5E |
| *dgk1Δ::leu1+ nem1Δ::kanR* | This study | SO9270 | S5C |
| *ept1-GFP::natR ade6-210 ura4-D18 leu1-32 h+* | This study | SO9297 | 6F |
| *ept1Δ::hphR ade6-210 ura4-D18 leu1-32 h+* | This study | SO9288 | 6G, 6I, S6C |
| *ept1Δ::hphR dgk1Δ::natR* | This study | SO9316 | 6H, 6I, S6D |
| *ept1Δ::hphR mCherry-atb2::hphR ura4+::cdc15<sup>S.p.</sup> promoter-NLS DG sensor (SV40 NLS-2x PKCβ DG binding domain-SV40 NLS)-GST-GFP* | This study | SO9312 | S6C |

Molecular genetic manipulations were performed using either a plasmid-based (Keeney and Boeke, 1994) or PCR-based (Bahler et al., 1998) homologous recombination. Lipid sensors were integrated into the *ura4* locus.

Overexpression of Dgk1 and its catalytic mutant were carried out by integration of pREP1-Dgk1 or the mutated version into the *leu1* locus of *S. pombe*. For overexpression experiments, cells were pre-grown in EMM containing 5µg/mL thiamine at 30°C for 18 hours. These cells were then collected by centrifugation and washed thrice in EMM. Cells were then diluted into separate flasks of EMM with or without thiamine and grown for 20 hours prior to imaging.

For experiments in Fig. 6A-E and Fig. S6A, B, cells were precultured in YES at 30°C for 18 hours. These were split into separate flasks and grown at 30°C for 18 hours in the appropriate medium (YES, EMM or EMM+Cho/Etn) prior to imaging the following day. Propidium iodide (Sigma-Aldrich) was used at a final concentration of 0.1µg/mL.

Growth rates were measured using a VICTOR Nivo Multimode Microplate Reader (PerkinElmer). Cells were initially precultured in YES at 30°C for 18 hours. These were diluted to OD_595_=0.05 and 200µL of each culture were seeded into 5 wells as technical repeats and grown at 30°C for 50 hours without shaking. Absorbance readings were obtained every 10min. Doubling time was calculated using the formula 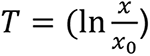, where T is the doubling time, x is the final OD_595_ and x0 is the initial OD_595_. Results were expressed as doubling time relative to the wild-type.

### Lipid sensor design

All lipid sensors were cloned into the pJK210-based plasmid backbone. DNA sequences of the lipid biosensors were synthesized using GeneWiz FragmentGENE service. *S. pombe* promoters (prom*^tdh1^*, prom*^cdc15^*) were inserted between KpnI-ApaI followed by the biosensor sequence between XhoI-EcoRI. This was immediately followed by the GST tag located between PacI-BamHI in the *S. pombe* sensors, followed by the fluorescent tag (GFP or mCherry) between BamHI-NotI with a stop codon located before the NotI restriction site (Fig. 1A). NLS version of the lipid biosensors contains two SV40 NLS motif appended to both ends of the lipid sensor module.

The PA lipid-binding module consisted of the *S. cerevisiae* Opi1 residues 111-189 to exclude the endogenous NLS signal located at residues 109-112, a duplication of the PA-binding amphipathic helices (residues 114-131) and a point mutation introduced at G120W (Fig. S1A) (Hofbauer et al., 2018). The DG biosensor consisted of the duplicated DG-binding domain of *R. norvegicus* PKCβ (residues 31-158) (Fig. S1C) (Lucic et al., 2016). PA-binding mutant was constructed with the following mutations: L124R, Y127A, L129R, M131A, I133R, K136A, K137A and R138A (Loewen et al., 2004), while DG-binding mutant contained point mutations at Q63E and Q128E (Lucic et al., 2016). The sequences of all sensor variants were codon-optimized for *S. pombe*.

### Lipidomics

Fission yeast cultures for lipidomics were grown in the defined EMM medium with all supplements (adenine, uracil, histidine and leucine) at 30°C. Cells were collected by filtration and snap-frozen in liquid nitrogen. Lipid extraction was performed using a two-phase chloroform-methanol extraction protocol (Ejsing et al., 2009). Cells were first disrupted in 200µL of water using a Beadruptor 12 (OMNI international) with ceramic beads at 4.75 speed setting for six cycles of 30s at 4°C. 100µL of the diluted lysate was transferred to fresh Eppendorf tubes and 900µL of chloroform-methanol 17:1 (v/v) was added. The mixture was left shaking at 4°C for 2h at 100rpm in an Eppendorf thermomixer shaker. Following that, the mixture was centrifuged at 9000rpm at 4°C for 2min. The lower phase was transferred to a fresh Eppendorf tube and dried in a Thermo Fisher Scientific Savant SpeedVac while a second chloroform-methanol 2:1 (v/v) extraction was performed with the upper phase. The second extraction was combined with the first and dried in the SpeedVac before resuspension in 200µL of chloroform-methanol 1:1 (v/v) containing internal standards (Table 2) and stored at -80°C until further use. Batch quality control samples were made by pooling and mixing 10µL of each sample lysate. These were aliquoted into 8 Eppendorf tubes for lipid extraction as described earlier. Blanks were prepared by using 100µL of water in place of lysates for lipid extraction. All lipid extracts from samples were mixed and aliquoted into 9 vials as technical quality control samples. The technical quality control was diluted with chloroform-methanol 1:1 (v/v) to prepare 100, 50, 25, 12.5 and 6.25% diluted samples to assess instrument response linearity.

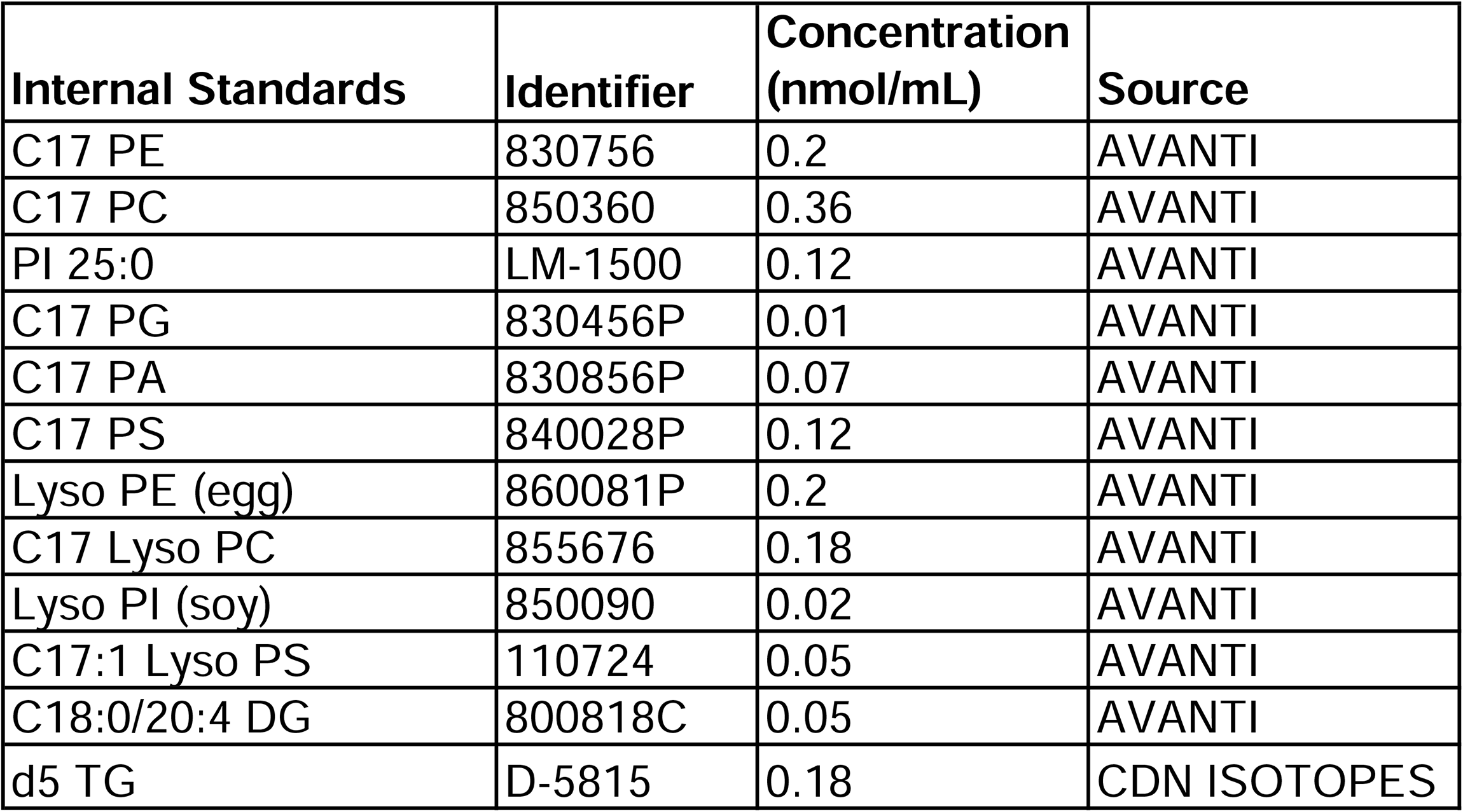

| **Internal Standards** | **Identifier** | **Concentration**<br>**(nmol/mL)** | **Source** |
| --- | --- | --- | --- |
| C17 PE | 830756 | 0.2 | AVANTI |
| C17 PC | 850360 | 0.36 | AVANTI |
| PI 25:0 | LM-1500 | 0.12 | AVANTI |
| C17 PG | 830456P | 0.01 | AVANTI |
| C17 PA | 830856P | 0.07 | AVANTI |
| C17 PS | 840028P | 0.12 | AVANTI |
| Lyso PE (egg) | 860081P | 0.2 | AVANTI |
| C17 Lyso PC | 855676 | 0.18 | AVANTI |
| Lyso PI (soy) | 850090 | 0.02 | AVANTI |
| C17:1 Lyso PS | 110724 | 0.05 | AVANTI |
| C18:0/20:4 DG | 800818C | 0.05 | AVANTI |
| d5 TG | D-5815 | 0.18 | CDN ISOTOPES |

Initial profiling of *S. pombe* lipids with isotopic correction were performed by Lipid Data Analyzer using the wild-type strain SO2865. Multiple reaction monitoring lists for each lipid class of interest were constructed from the data obtained from the initial profiling experiments and used for targeted LC-MS lipidomics (Tables 3 and 4). All LC-MS/MS was performed on an Agilent 6495A QqQ mass spectrometer connected to a 1290 series chromatographic system with electrospray ionization for lipid ionisation. All sample injection volumes were set at 2µL. The spray voltage and nozzle voltage were set at 3500V and 500V respectively. The drying gas and sheath gas temperatures were maintained at 200°C and 250°C respectively, with flow rates both set at 12L/min. The nebulizer setting was 25psi. Following instrument stabilisation with 15 injections of a QC sample, instrument stability was monitored by an injection of a QC sample and blank sample every 5 sample injections.

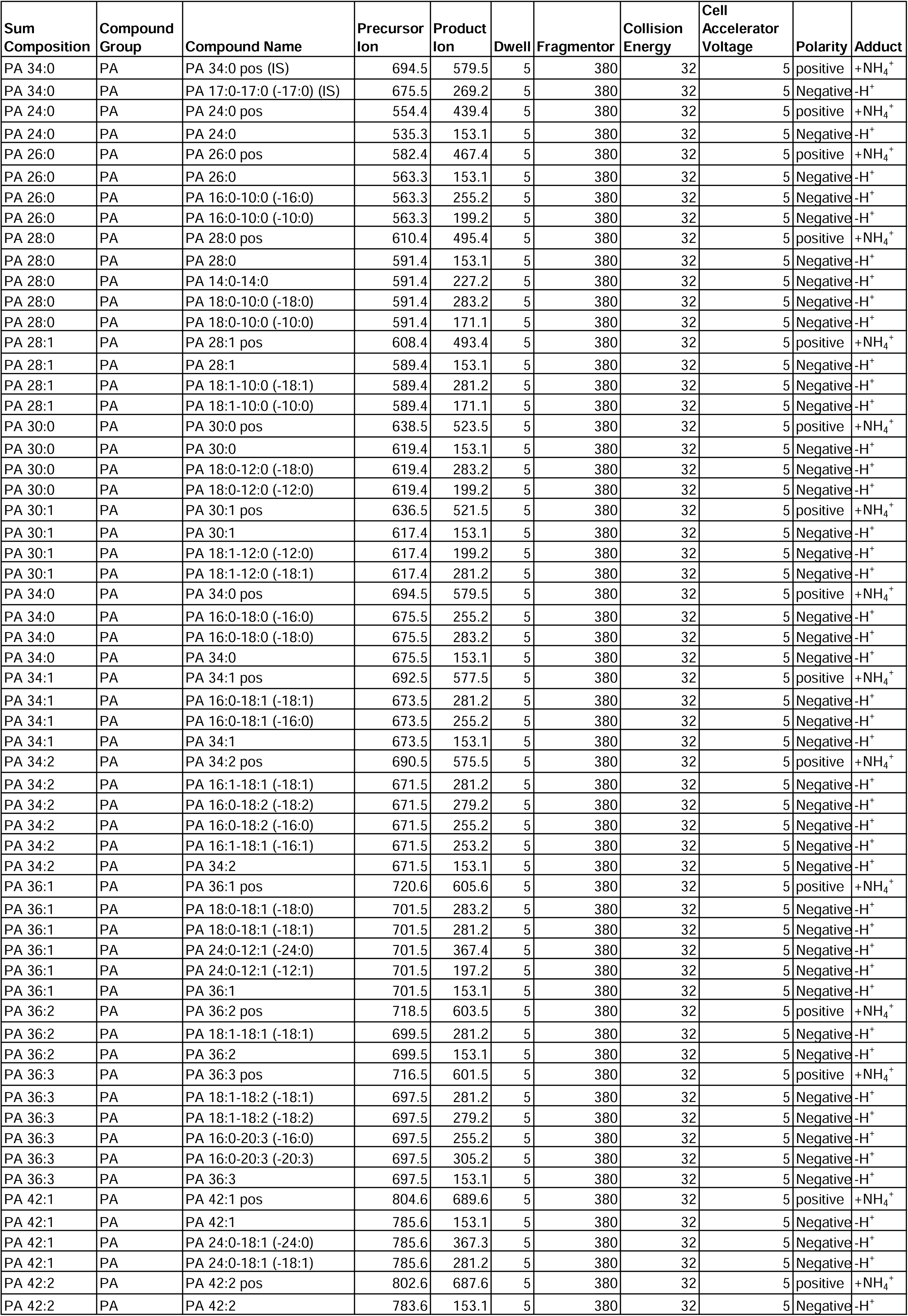
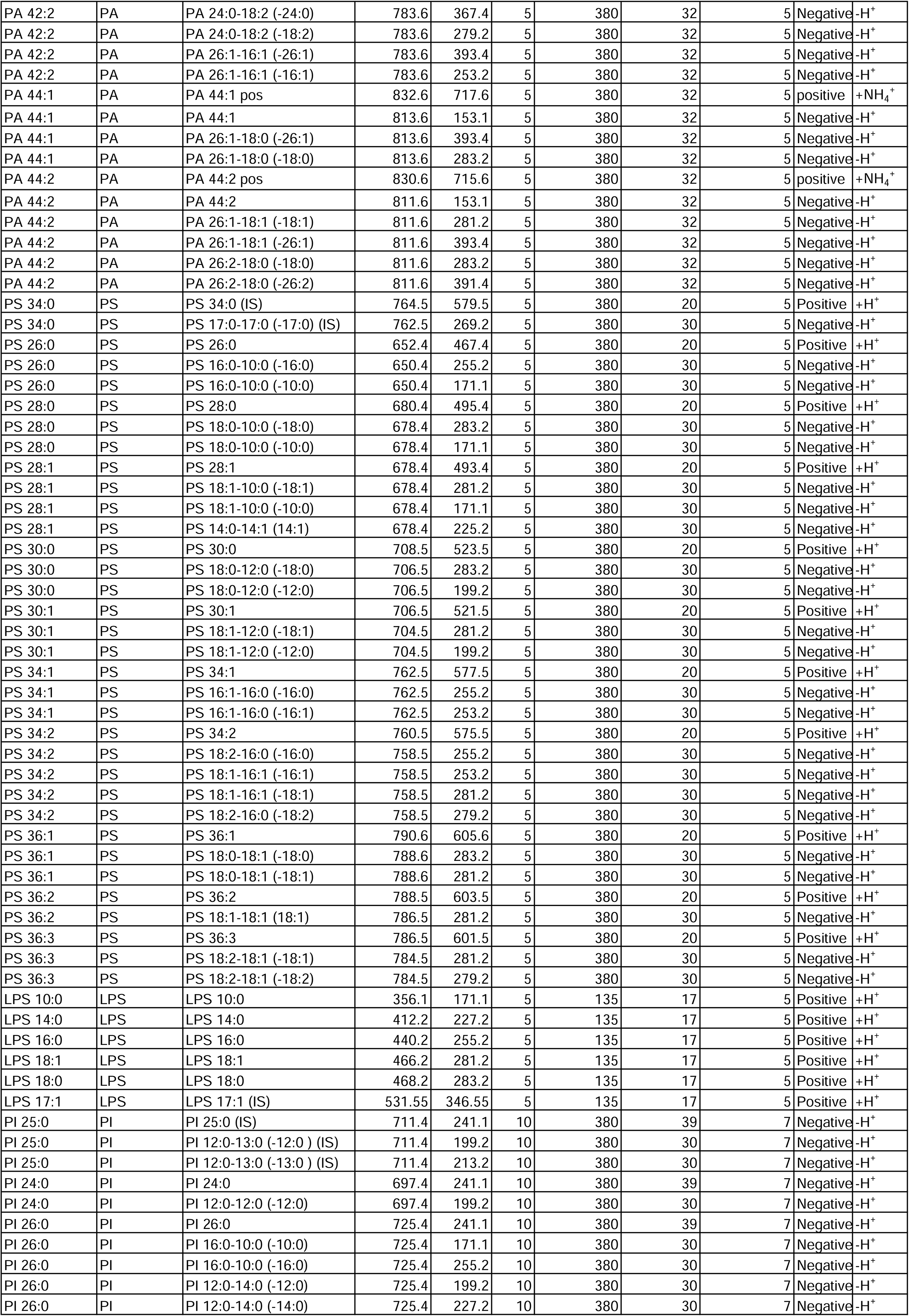
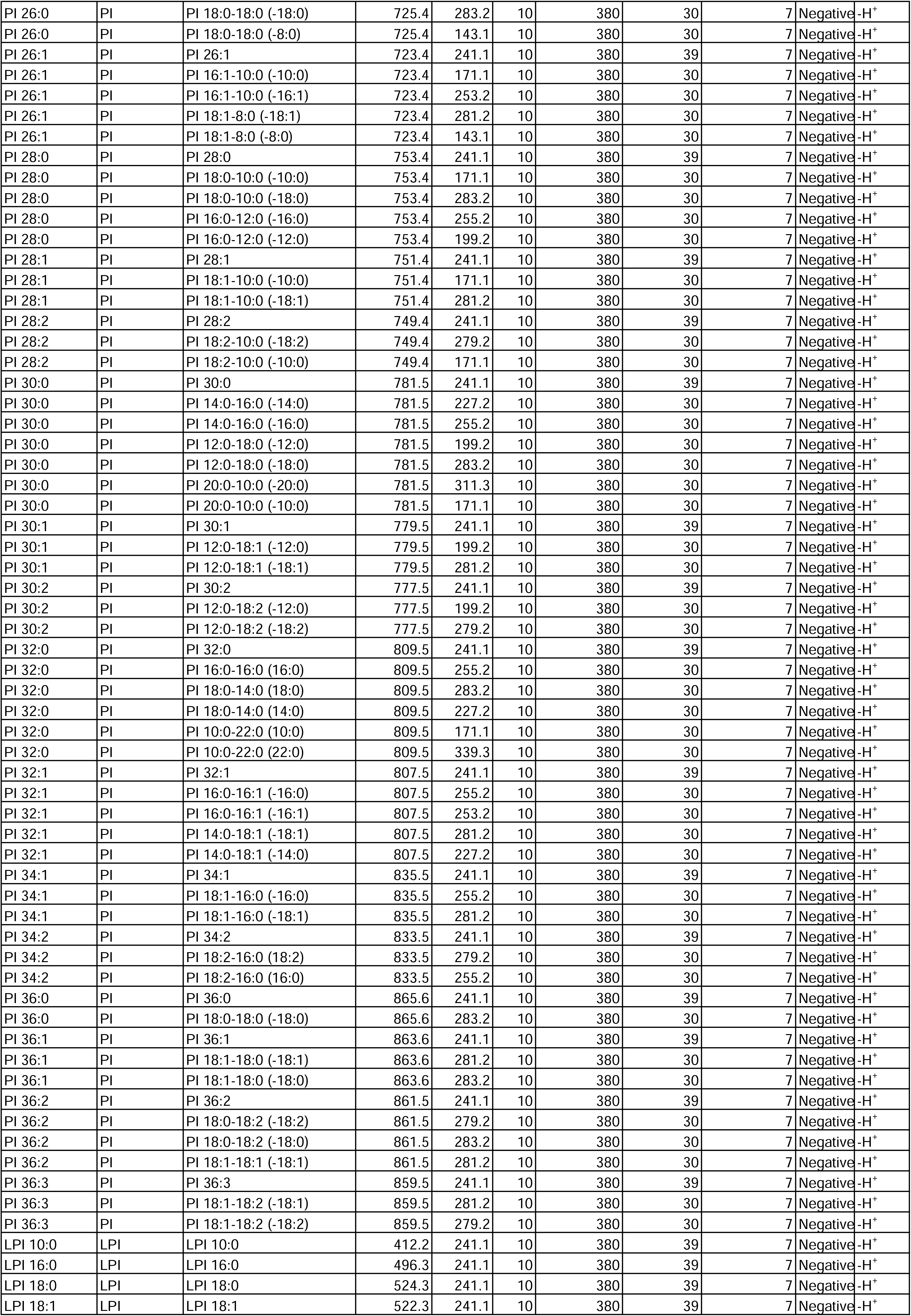
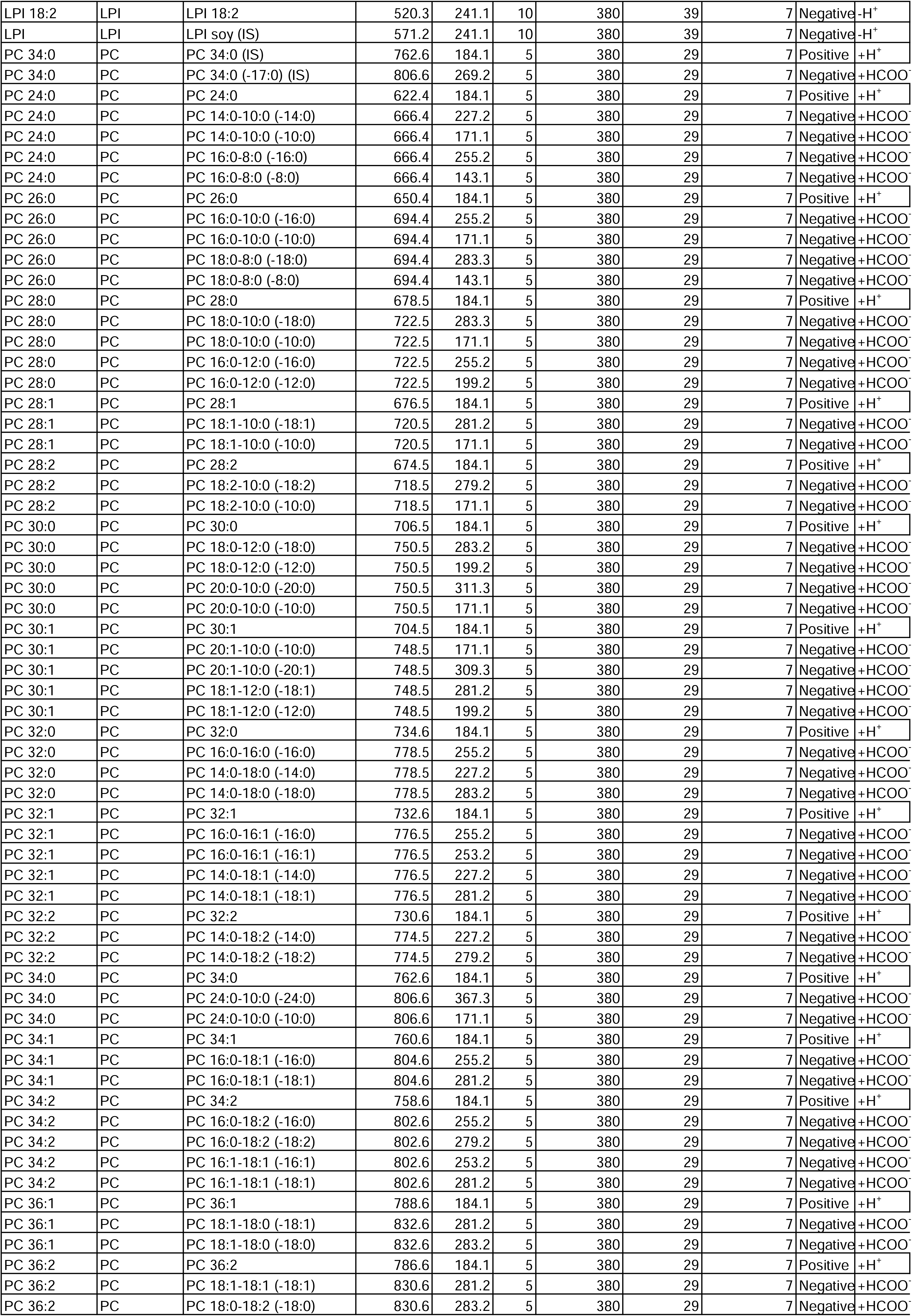
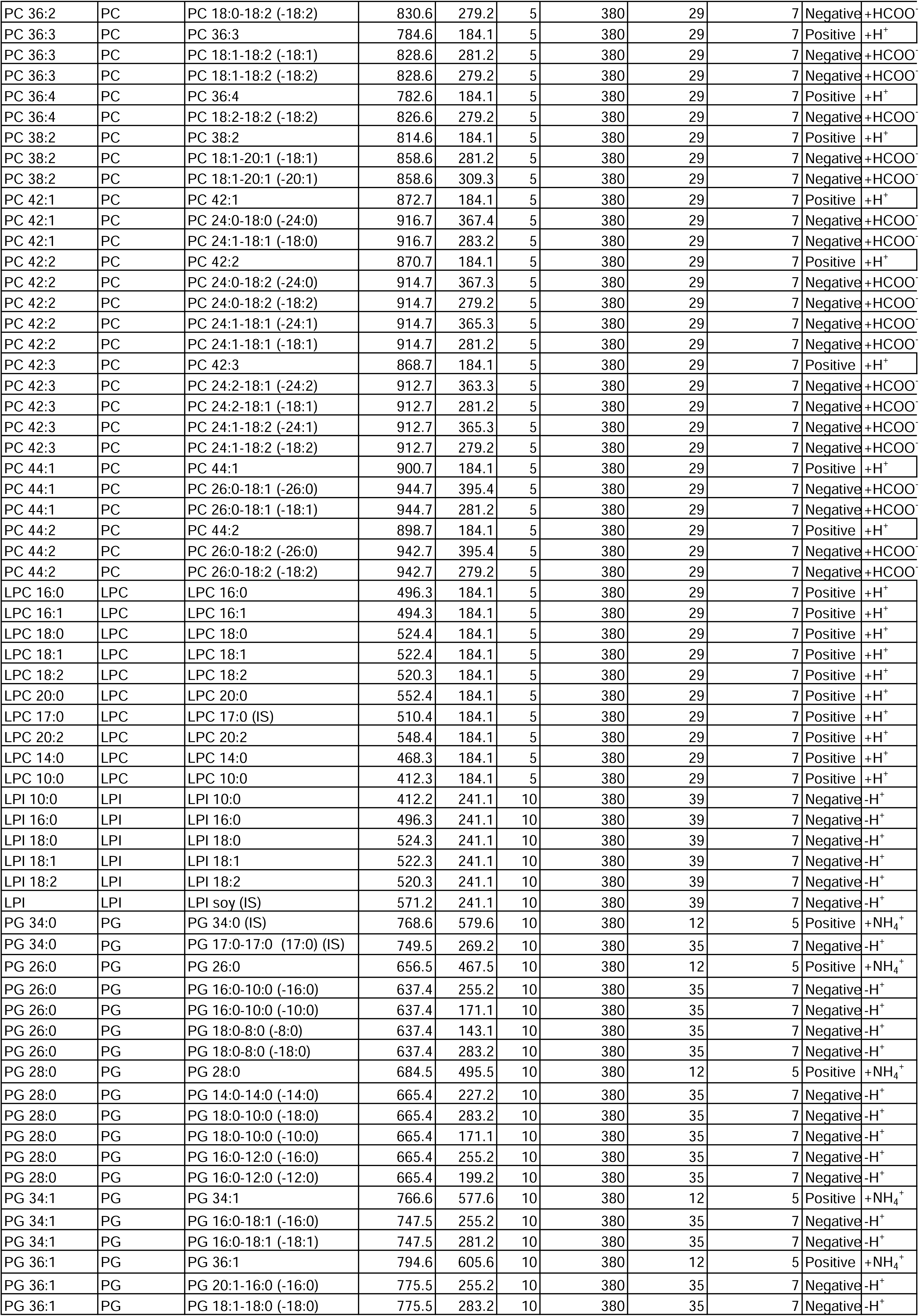
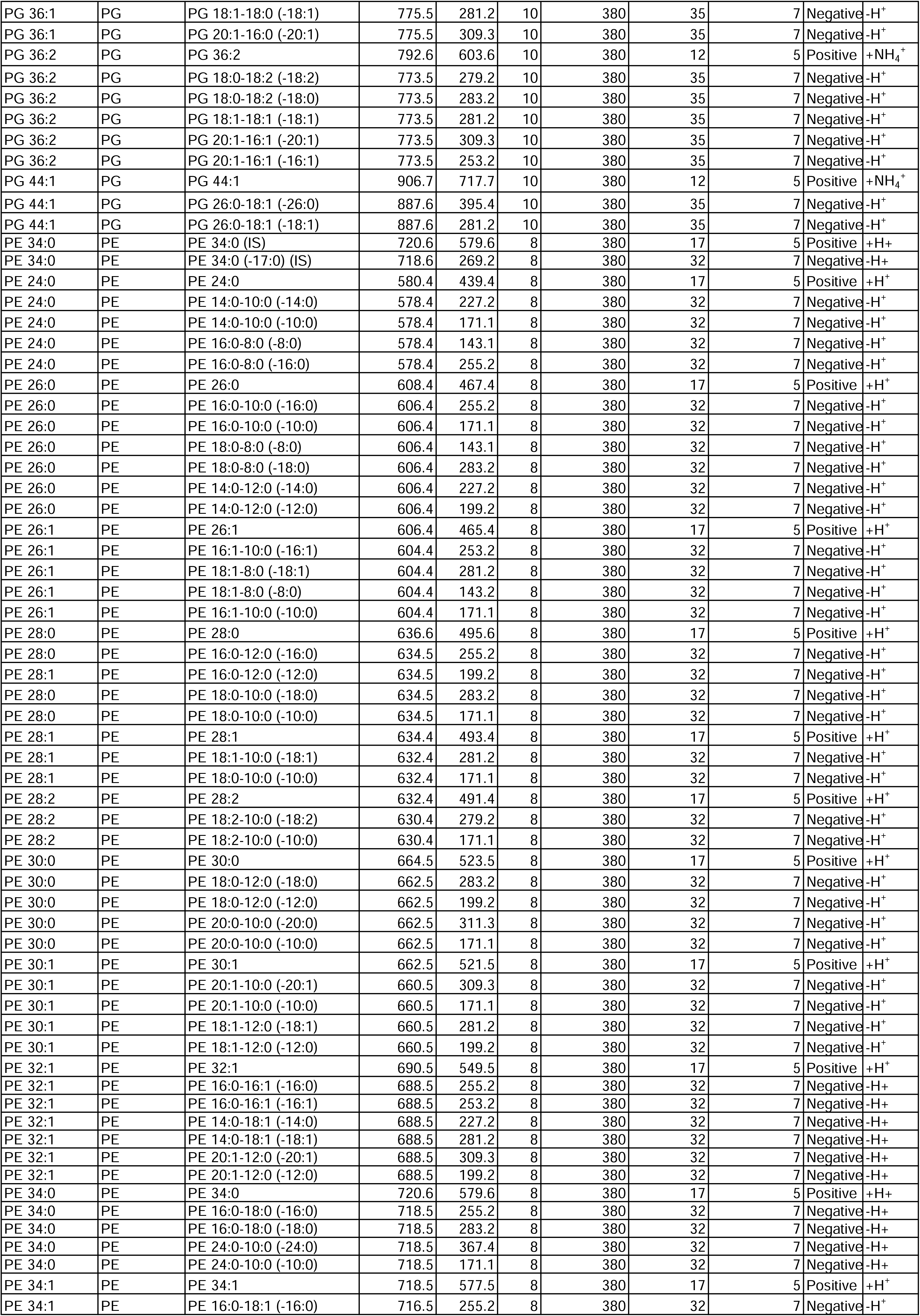
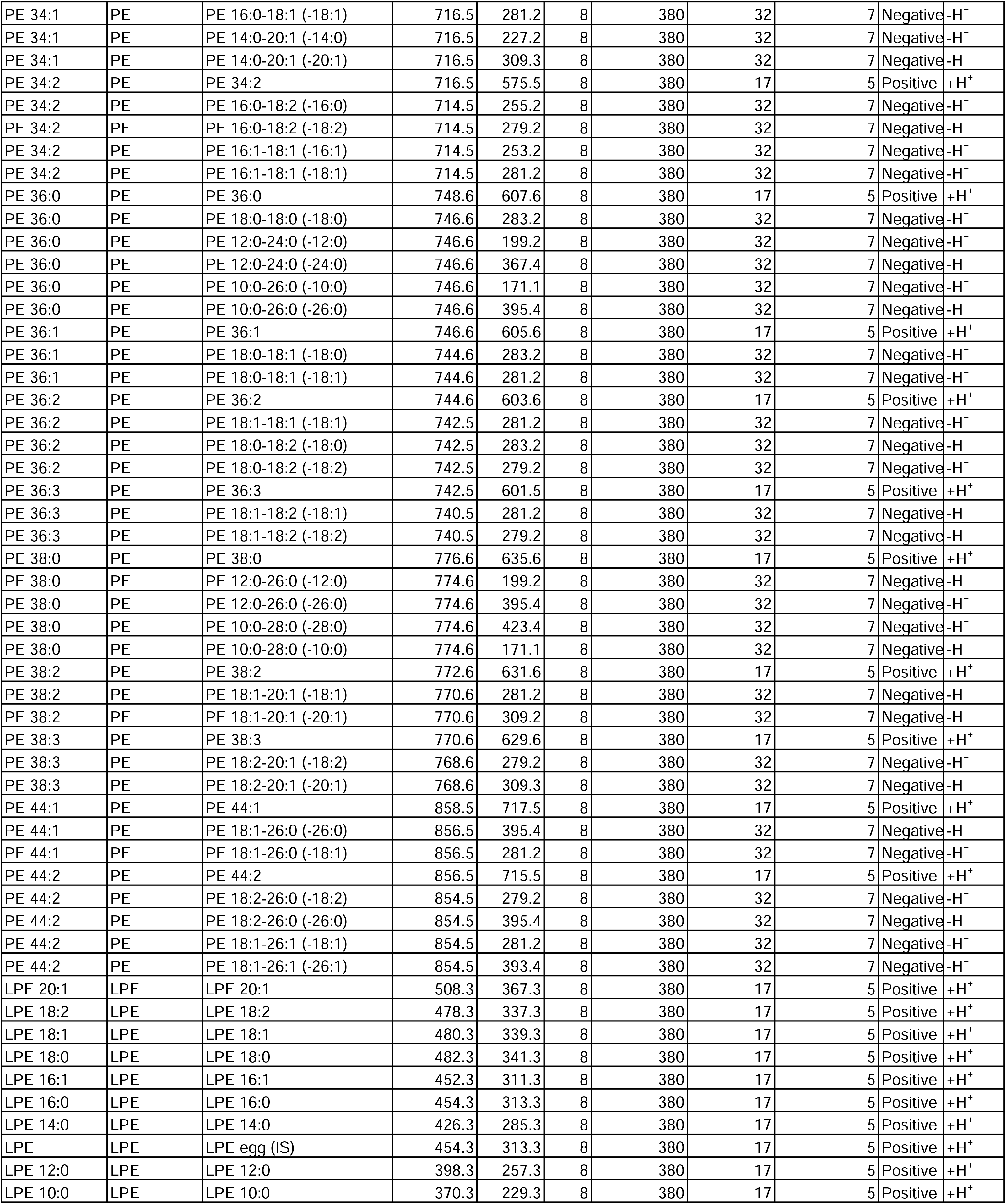

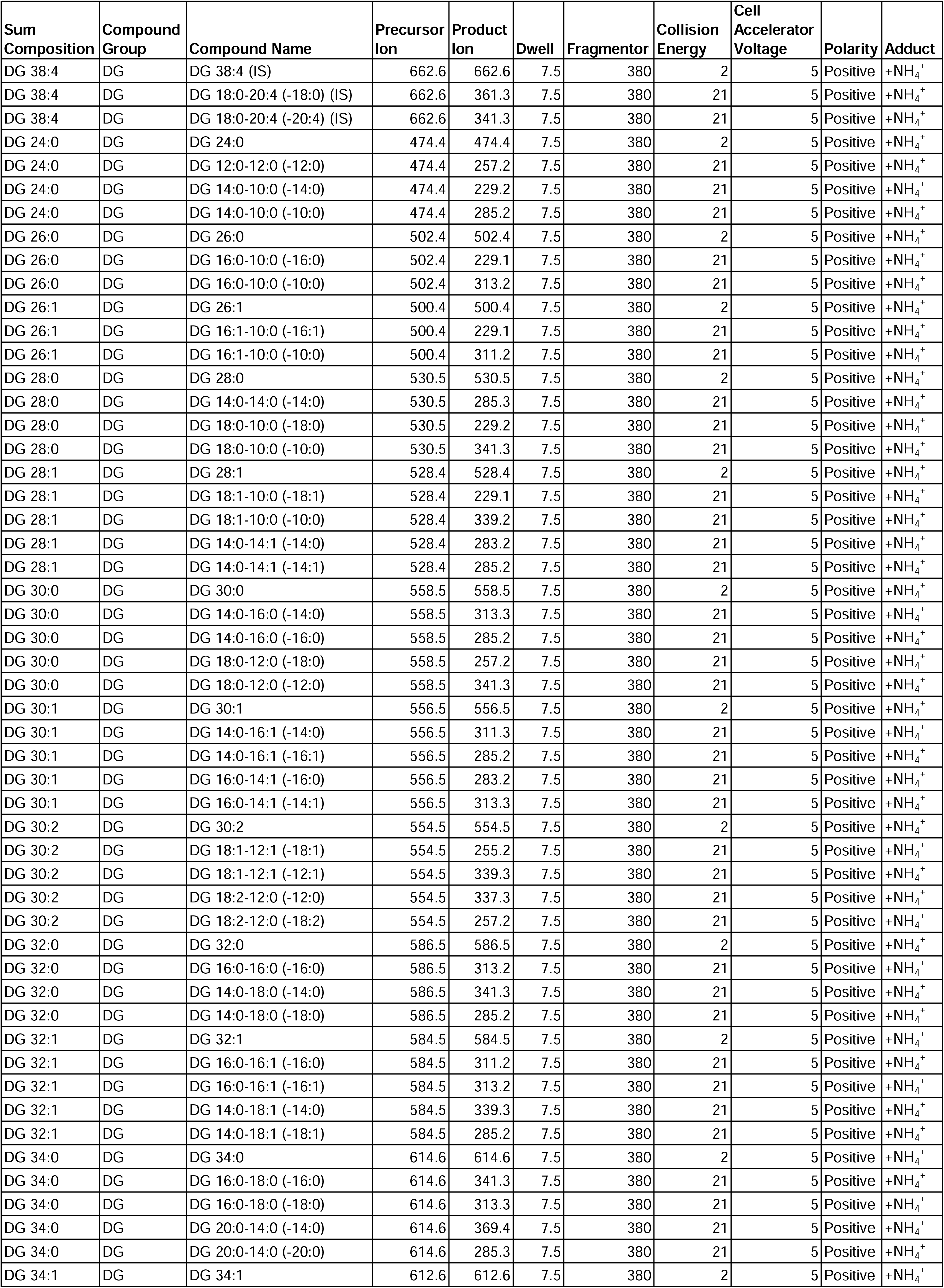
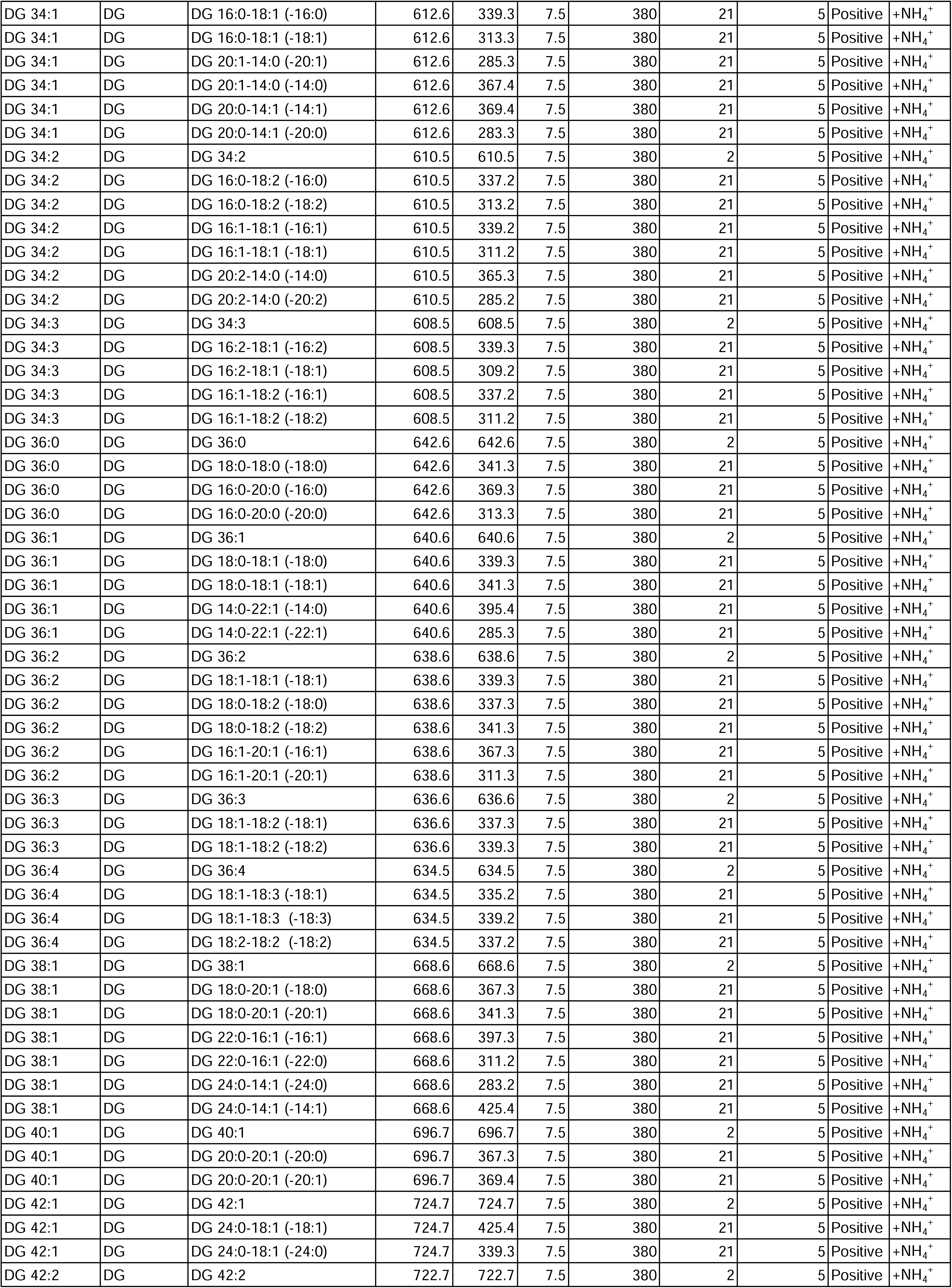
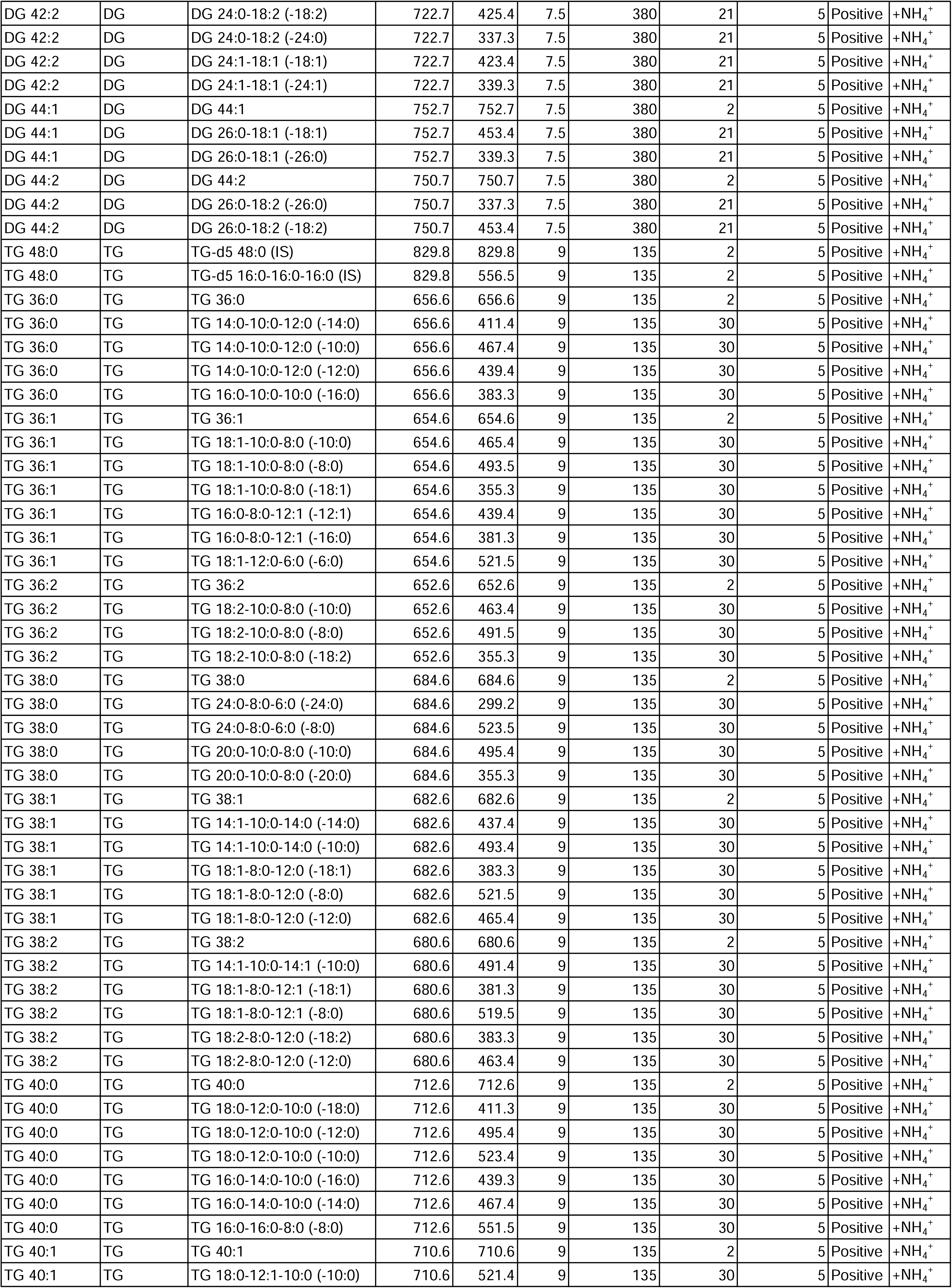
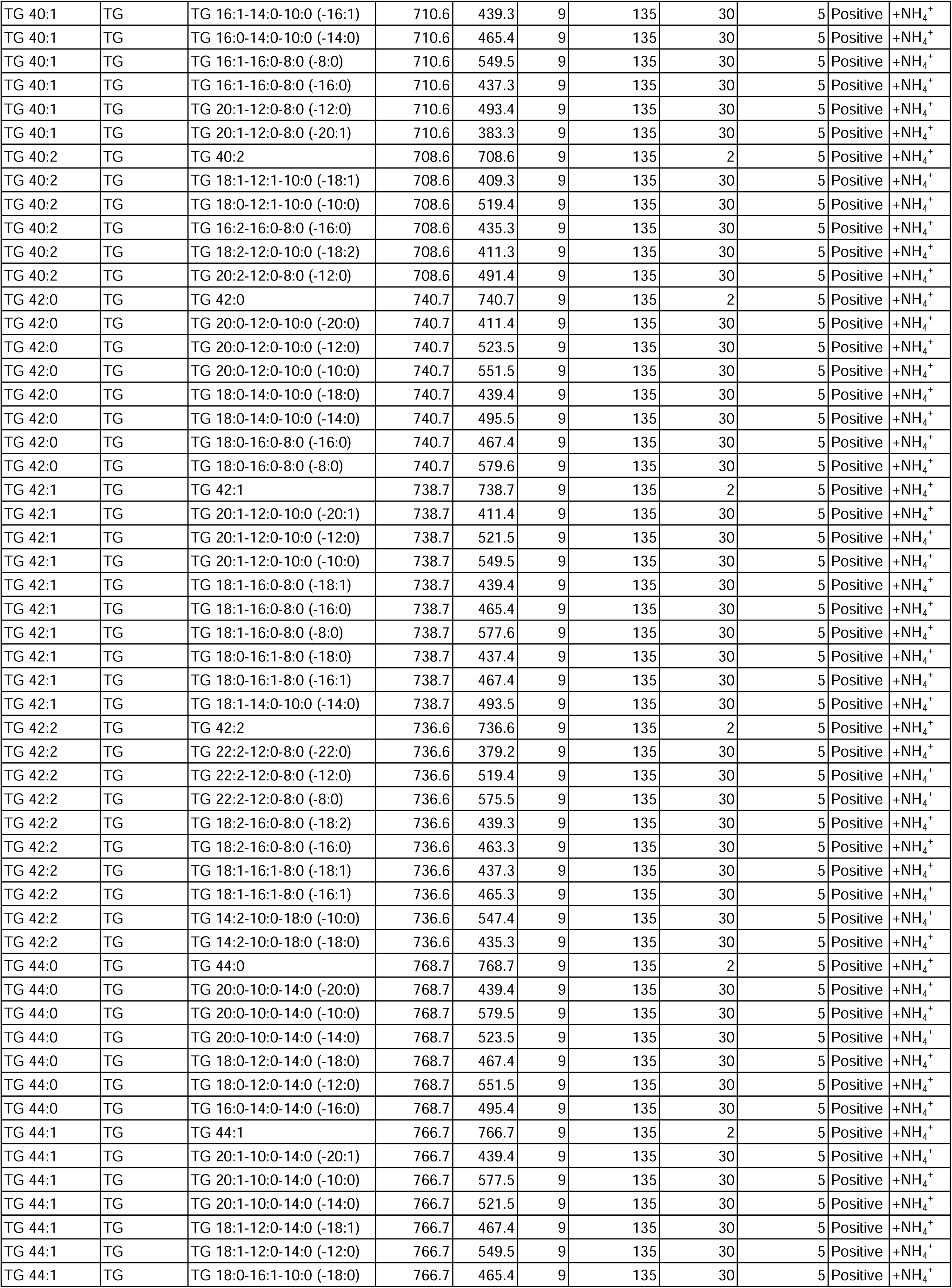
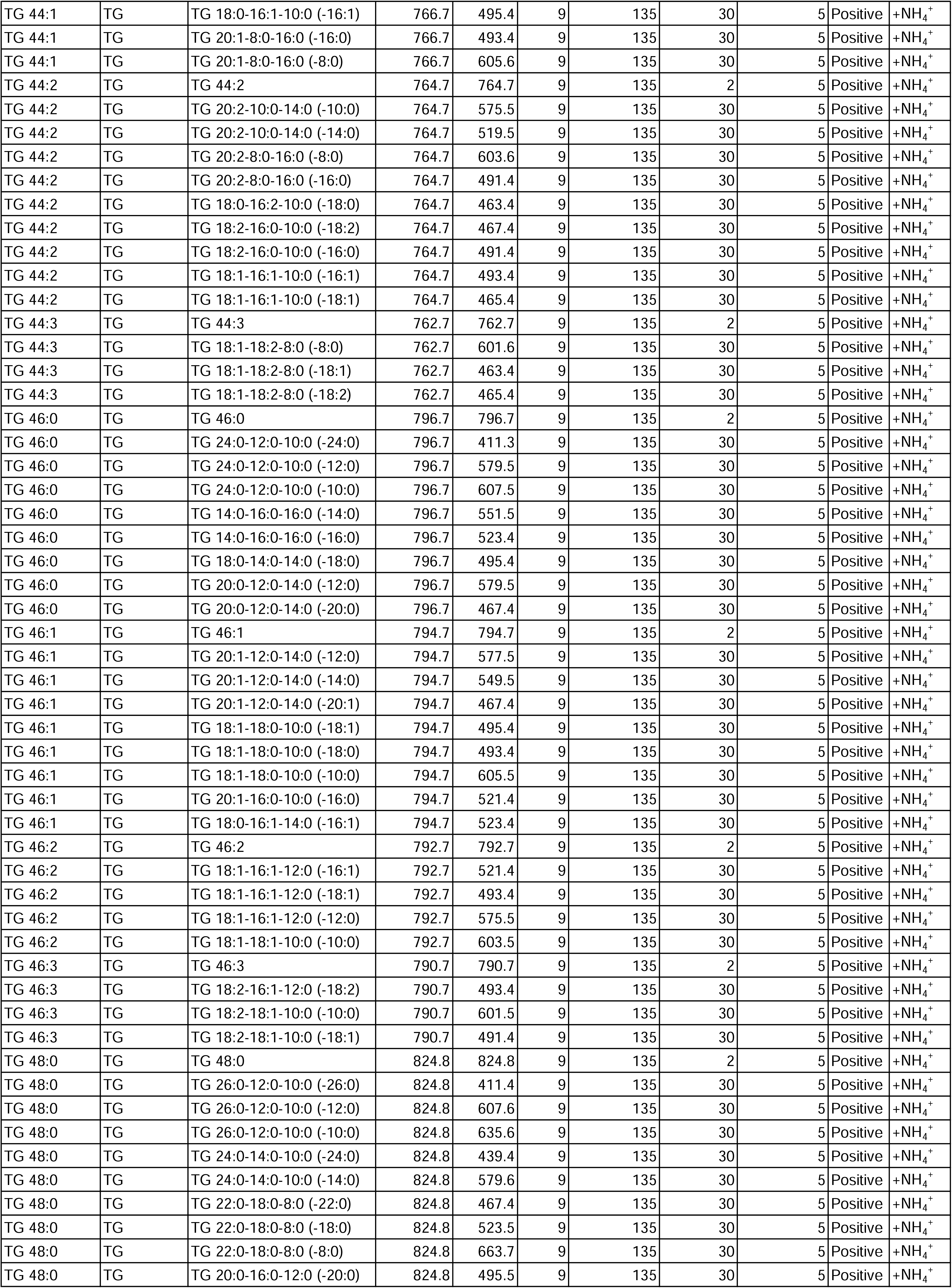
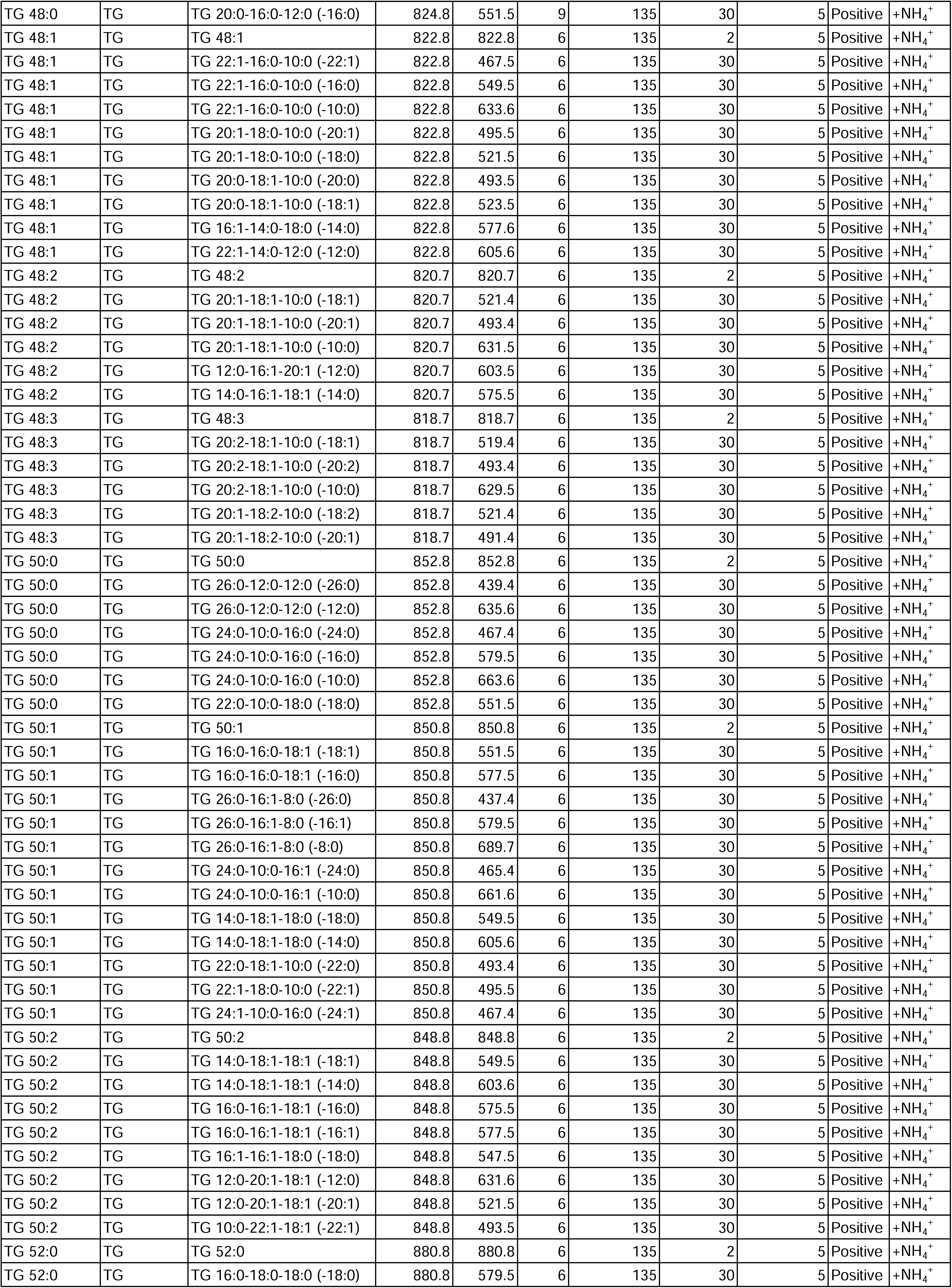
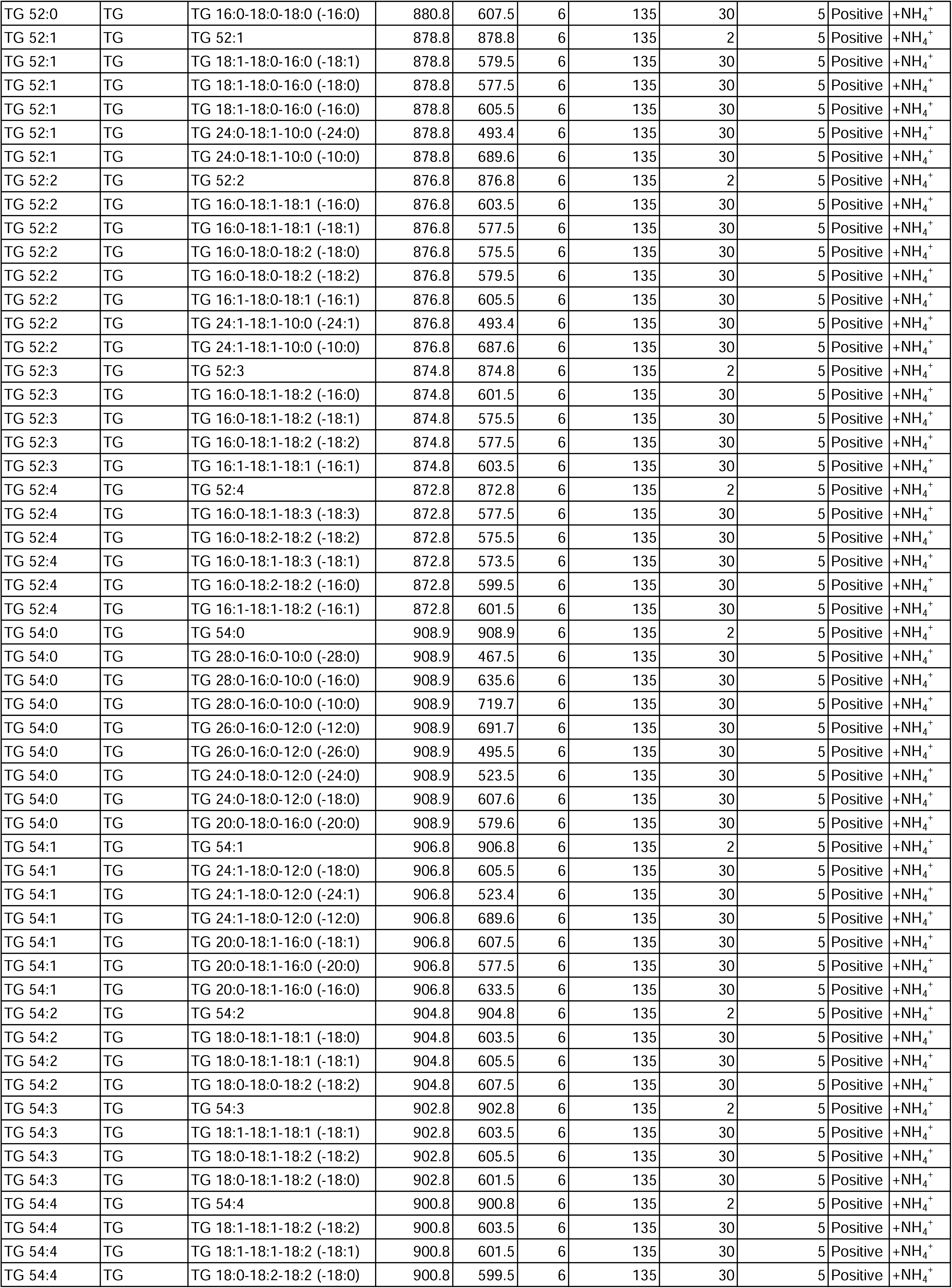
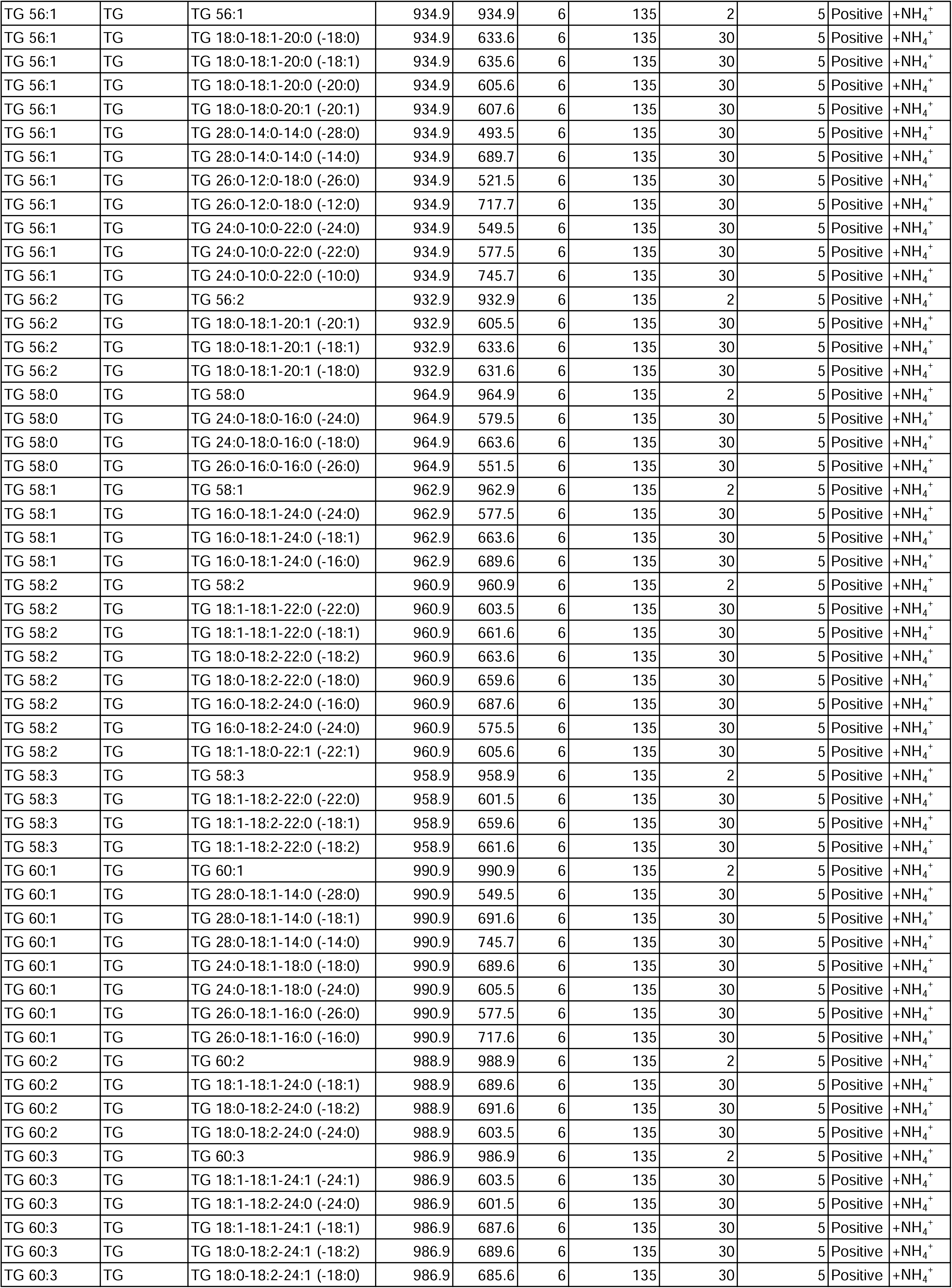

Chromatographic separation of the GPLs phosphatidylinositol, phosphatidylcholine, phosphatidylethanolamine and phosphatidylglycerol was performed using a gradient elution on an ethylene bridged hybrid column with solvent A 1:1 (v/v) acetonitrile/ H_2_O with 10mM ammonium formate, and solvent B containing 19:1 (v/v) acetonitrile/ H_2_O with 10mM ammonium formate. The flow rate was set at 0.4mL/min with solvent B set at 40% upon injection and increasing to 100% in 7min. This value was retained for 2min, decreased back to 40% in one minute, and then retained there until the end of the gradient by 14min. The eluent was directed to the electron spray ionization source of the mass spectrometer operated in positive ionisation.

Chromatographic separation of PA and phosphatidylserine was performed using a different liquid phase (Triebl et al., 2014). Mobile phase A was deionized water containing 10mM ammonium formate and 0.5% formic acid. Mobile phase B was 2-propanol/acetonitrile 5:2 (v/v) containing 10mM ammonium formate and 0.5% formic acid. Gradient elution began at 5% solvent A with a linear increase to 50% over 12 min; the 50% solvent A was held for 3min, and lastly, the column was re-equilibrated for 15min.

Chromatographic separation of neutral lipids DG and TG was performed using reversed-phase liquid chromatography on an Agilent rapid resolution HD Zorbax Eclipse-C18 column. The mobile phases A consisted of 3:2 (v/v) H_2_O/ acetonitrile with 10mmol/L ammonium formate and B consisted of 1:9 (v/v) acetonitrile/ isopropanol with 10mmol/L ammonium formate. The flow rate was set at 0.4mL/min with solvent B set at 20% upon injection and increasing to 60% in 2min, increasing to 100% over 12min and held there for a further 2min. This was then decreased to 20% for the next 2min.

Raw data were processed with MassHunter QqQ Quantitative software (version B.08). Areas under the curve of the chromatogram peaks for each transition were measured and exported to Excel. Normalised peak areas were calculated by dividing the peak areas of the analyte with the corresponding internal standard. Relative abundances were obtained by multiplying the normalised peak areas with the molar concentration of the corresponding internal standard.

All lipidomics experiments were performed with three biological replicates and results presented as mean ±SD. Statistical analysis was performed using non-parametric heteroscedastic t-test. All solvents used were LC-MS grade purchased from Sigma-Aldrich or Thermo Fisher Scientific. Annotation of lipid classes and species was done according to the classification system previously described (Liebisch et al., 2013).

### Fluorescence-activated cell sorting

*S. pombe* cells were grown at 30°C in YES medium and 10^7^ cells were collected. Cultures were diluted every 24 hours to OD_595_=0.0005, and samples at OD_595_=0.5 were typically collected over the course of three days. Cell pellets were resuspended in 1mL of 70% cold ethanol and kept at 4°C until required. For staining of cells for FACS analysis, cells were first centrifuged at 6000rpm for 2min at 4°C and the supernatant removed. The pellet was washed once in 1mL of 50mM sodium citrate and resuspended in 500µL of 50mM sodium citrate containing 0.1mg/mL RNaseA (Thermo Fisher Scientific). This was incubated at 37°C for 18 hours with gentle shaking. 500µL of 50mM sodium citrate containing either 2µM SYTOX green (Thermo Fisher Scientific) for *S. pombe* samples or 8µg/mL propidium iodide (Sigma-Aldrich) for *S. japonicus* samples was added, and the suspension sonicated briefly to remove cell doublets before FACS analysis. All FACS experiments were performed on a LSRFortessa Cell Analyzer (BD Biosciences) at the Crick Flow Cytometry Science Technology Platform, with at least three biological replicates.

### Quantification and statistical analyses

Image processing and quantifications were performed in Fiji (Schindelin et al., 2012). Circularity of the nucleus was calculated using the formula 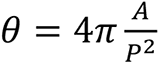 where θ is circularity, A is the area and P is the perimeter.

Quantification of PA and DG at the INM or cortex of mitotic cells were obtained using the segmented line function in Fiji with a line width of 5 pixels every 1 minute of the time-lapse images. Lipid sensor decay at the NE of mitotic cells was normalized by taking the ratio of the sensor intensity at the NE to the cortex. The lines and shaded area represent the mean ±SD respectively. The rate of sensor decay at the NE was calculated by taking the gradient via linear regression of the sensor decay at the NE in individual cells. For quantification of PA and DG at the INM of interphase cells (Fig. 1H, 1I, 2D, 2F, 5D, 5E and 6E), segmented line function in Fiji with a line width of 5 pixels was used. To account for variation in our sensor expression levels, average whole cell fluorescence intensity was also measured using the polygon function in Fiji, and values above 1.5 interquartile range were discarded.

Box-and-whiskers plots were created using the Tukey method in Prism 7 (GraphPad Software). All statistical analyses were performed using non-parametric heteroscedastic t-test.

### Image acquisition

Working concentration of all BODIPY-based dyes (Thermo Fisher Scientific) used was 1µM. Time-lapse microscopy of cells undergoing mitosis was performed using cells grown on agar pads (Pemberton, 2014). All images were obtained using a Yokogawa CSU-X1 spinning disk confocal system mounted on the Eclipse Ti-E Inverted microscope with Nikon CFI Plan Apo Lambda 100X Oil N.A. = 1.45 oil objective, 600 series SS 488nm, SS 561nm lasers and Andor iXon Ultra U3-888-BV monochrome EMCCD camera controlled by Andor IQ3 or Fusion software. Temperature was maintained at 30°C, unless otherwise specified, using an Oko Cage Incubator. Maximum intensity projections of 3 Z-slices of 0.5μm step size images are shown in time-lapse montages.

## Acknowledgements

We are grateful to the Oliferenko lab for discussions and Eugene Makeyev for suggestions on the manuscript. Many thanks to Lydia Thompson and the Crick Flow Cytometry STP for help with FACS experiments; Foo Juat Chin and the Singapore Lipidomics Incubator lab members for help with lipidomics experiments. Sherman Foo was in part supported by the King’s-NUS Joint PhD scholarship. Work in M.R.W. laboratory is supported by grants from the National University of Singapore via the Life Sciences Institute, the National Research Foundation (NRFSBP-P4) and the NRF and A*STAR IAF-ICP I1901E0040. Work in S.O. lab was supported by the Francis Crick Institute, which receives its core funding from Cancer Research UK (FC001002), the UK Medical Research Council (FC001002), and the Wellcome Trust (FC001002), and the Wellcome Trust Senior Investigator Award (103741/Z/14/Z), Wellcome Trust Investigator Award in Science (220790/Z/20/Z) and BBSRC (BB/T000481/1) to Snezhana Oliferenko.

## Author contributions

Sherman Foo conceived and performed cell biological and biochemical experiments; generated strains; analyzed data; and co-wrote the manuscript. Sherman Foo, Amaury Cazenave-Gassiot and Markus R. Wenk designed, performed, and interpreted all lipidomics experiments. Snezhana Oliferenko conceived and interpreted experiments, co-wrote and edited the manuscript.

## Supplemental figure legends

**Figure S1.**
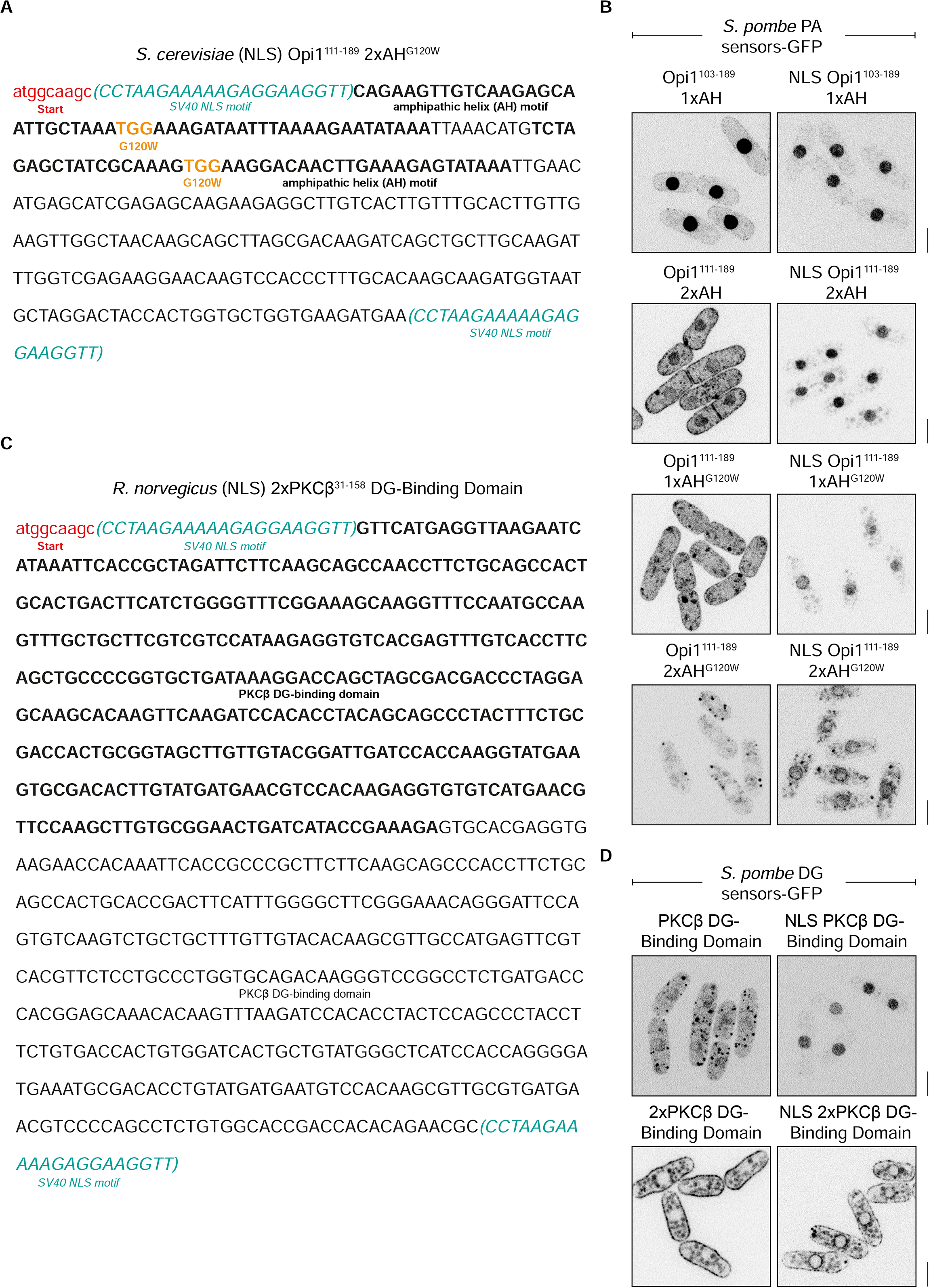
Development and optimization of lipid biosensors in fission yeast. (**A**) Sequence of the optimized PA lipid sensor module. The optimized PA lipid sensor consists of two consecutive *S. cerevisiae* Opi1 amphipathic helices each containing a G120W point mutation. The amphipathic helices are indicated in bold, mutations indicated in orange, the initiation codon and linker Ala-Ser in red, and SV40 NLS motifs in cyan. (**B**) These single-plane spinning-disk confocal images show optimization of the PA lipid sensors. The single Opi1 amphipathic helix contains an endogenous NLS at residues 109-112. Removal of this endogenous NLS and doubling of the amphipathic helix improves membrane enrichment. Introduction of a single G120W mutation predicted to improve its PA specificity leads to an increased association of the sensors to LDs and the INM. The final iteration of the PA sensors used throughout this work includes two amphipathic helices and the G120W mutation. (**C**) Sequence of the optimized DG lipid sensor module. The optimized DG lipid sensor consists of two consecutive *R. norvegicus* PKCβ DG-binding domains. The first domain is indicated in bold, the initiation codon and linker Ala-Ser in red, and SV40 NLS motifs in cyan. (**D**) Doubling the PKCβ DG-binding domain leads to membrane enrichment of the DG sensors. (**C**, **D**) Scale bars represent 5µm.

**Figure S2.**
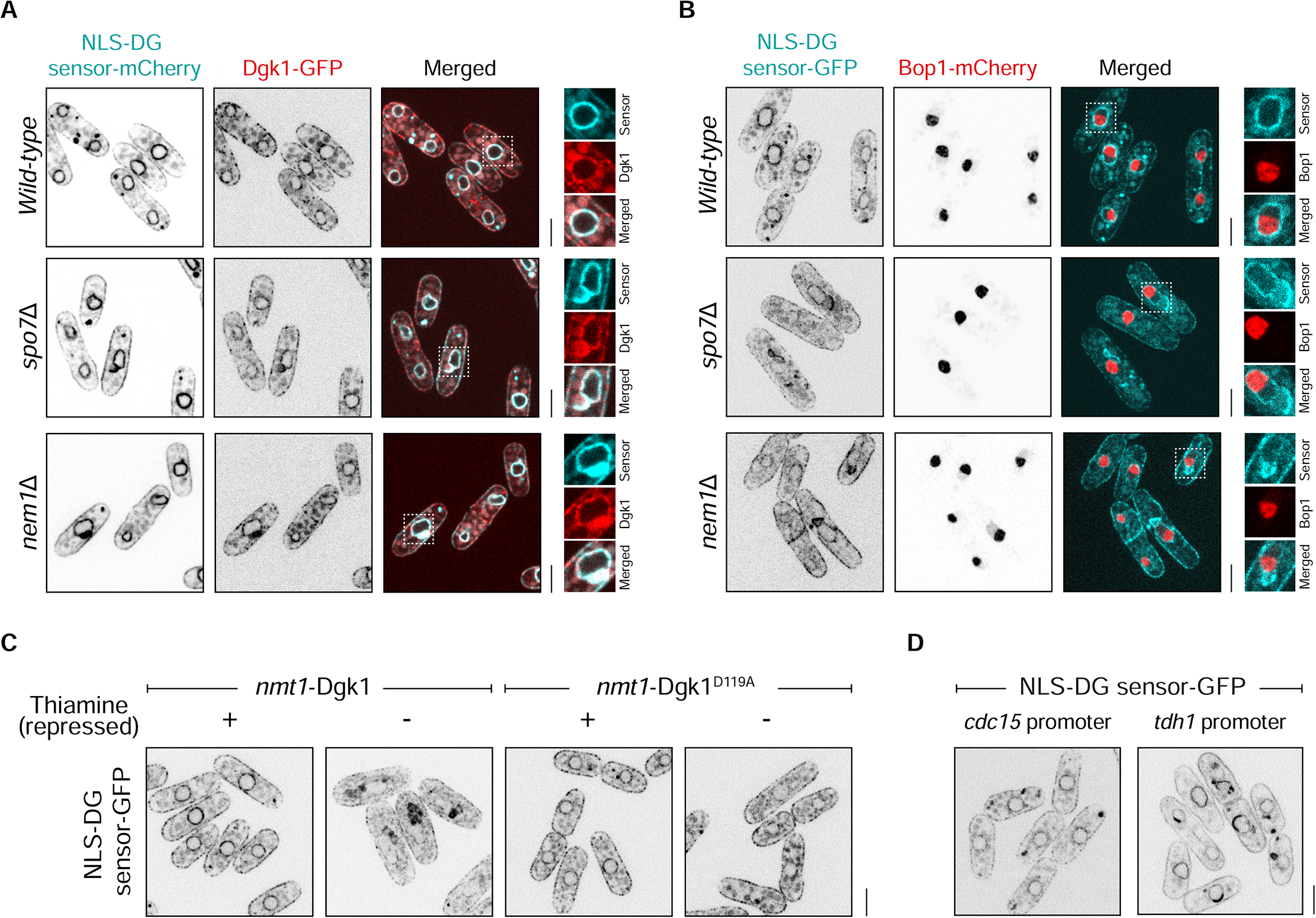
Excess NLS-DG sensor may lead to its clustering at the INM. (**A**) These single-plane spinning-disk confocal images show that the DG kinase Dgk1-GFP is not excluded from bright NLS-DG-mCherry patches at the INM in *spo7*Δ and *nem1*Δ cells. (**B**) Bright NLS-DG-GFP patches at the INM in YES-grown *spo7*Δ and *nem1*Δ cells are not associated with the nucleolus marked by Bop1-mCherry. These bright patches of NLS-DG-GFP sensor can be also induced by overexpression of Dgk1 causing a drop in DG (**C**) or an increase in the absolute amount of the sensor in a cell by expressing it from a stronger *tdh1* promoter (**D**). The wild-type Dgk1 and the catalytically inactive D119A mutant were expressed under the control of the thiamine-repressible *nmt1* promoter. (**A**, **B**) Insets represent magnified areas indicated by the dotted lines. (**A**-**D**) Scale bars represent 5µm.

**Figure S3.**
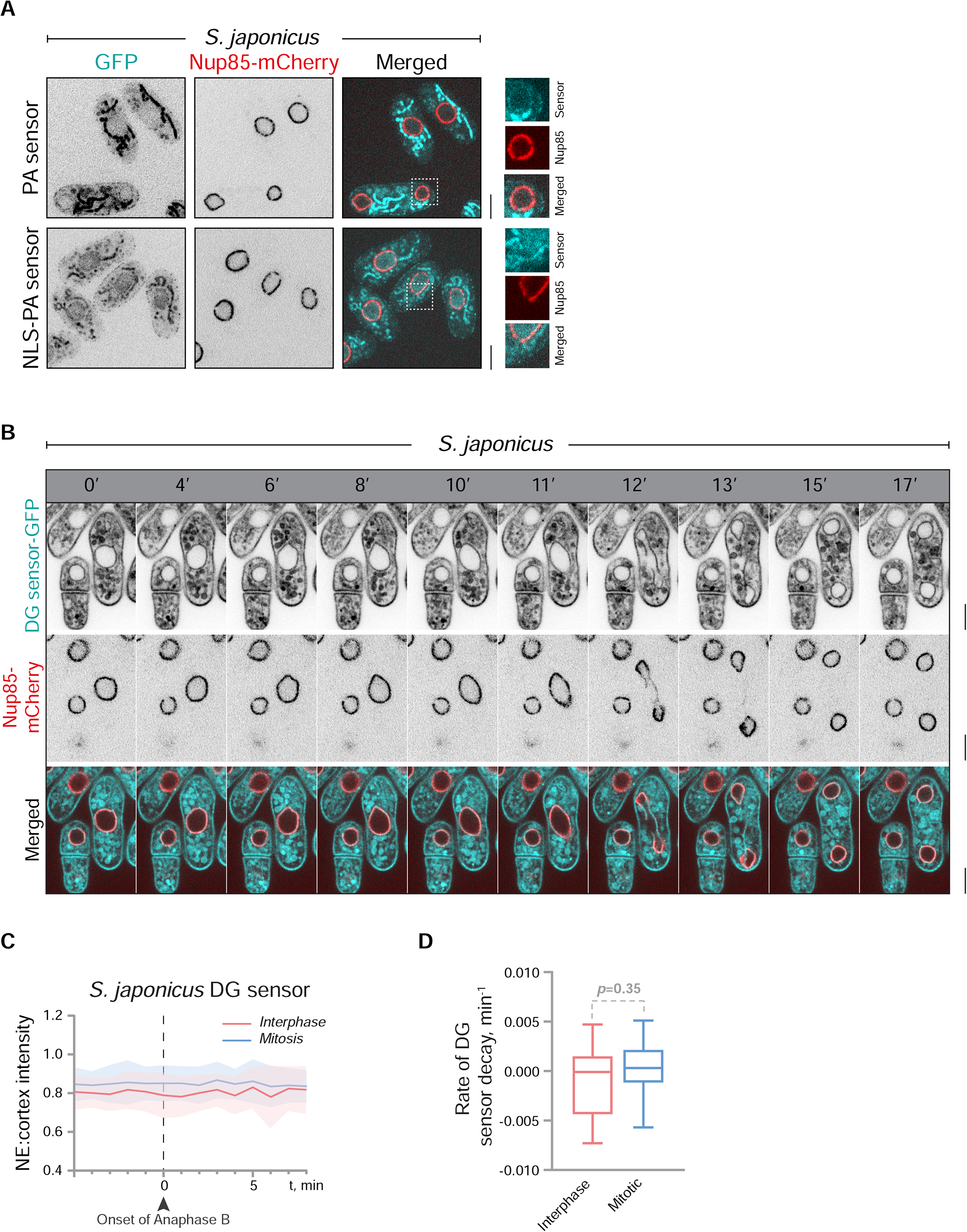
The subcellular distribution of PA and DG in *S. japonicus* differs significantly from *S. pombe*. (**A**) Wild-type *S. japonicus*, unlike *S. pombe*, shows a very weak PA enrichment at the INM. (**B**) In addition to the INM, DG is also enriched at the ONM of *S. japonicus*, as shown in this spinning disk time-lapse microscopy montage of *S. japonicus* expressing DG sensor. Shown are the maximum projections of three Z-slices. (**C**) The DG sensor intensity at the ONM remains relatively constant throughout the semi-open mitosis of *S. japonicus*, quantified in (**D**). (**A**, **B**) Nucleoporin Nup85-mCherry marks the nuclear boundary; scale bars represent 5µm. (**C**, **D**) n=10 cells, p-values are derived from the unpaired t-test.

**Figure S4.**
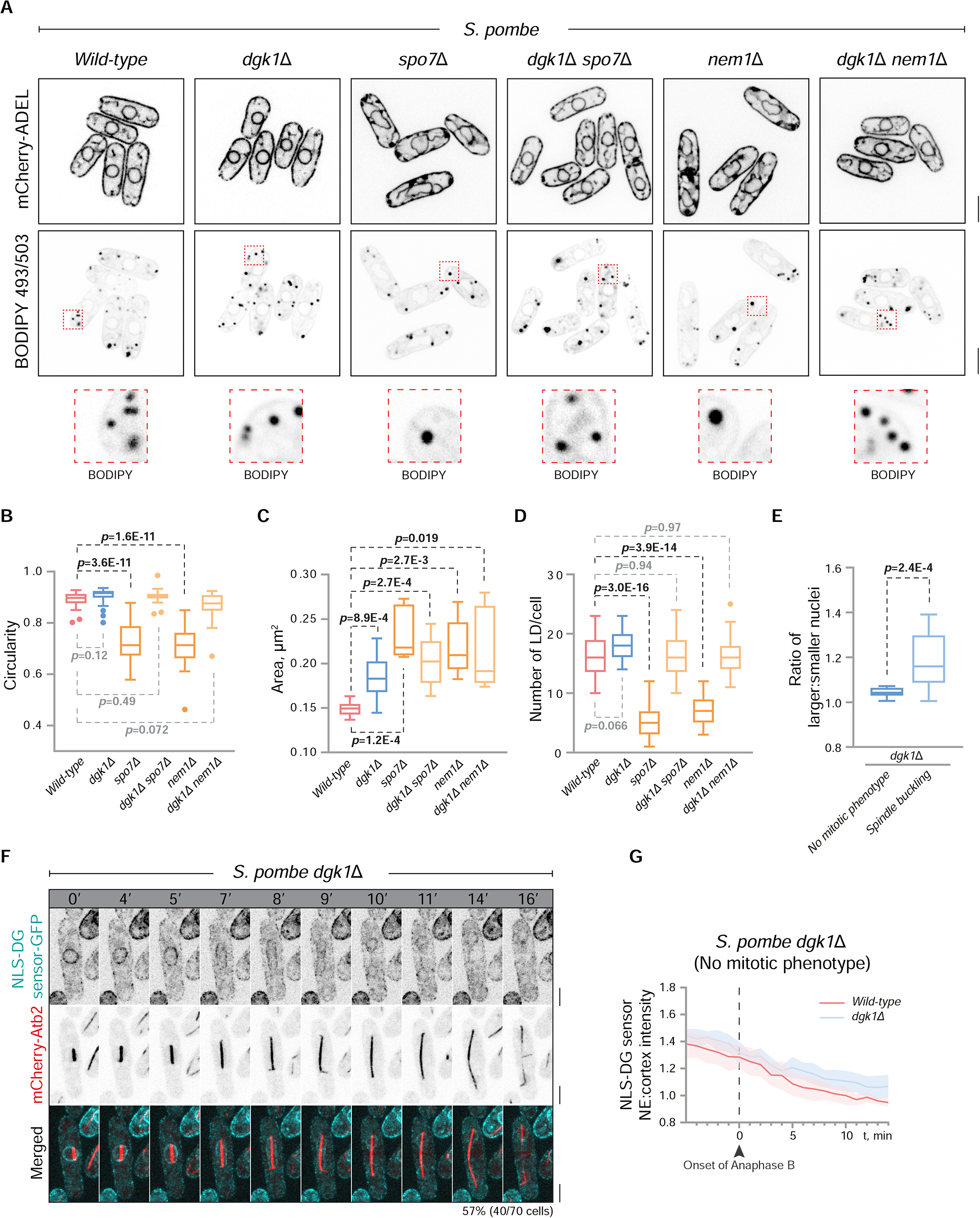
Dgk1 rescues the NE expansion phenotype of the lipin pathway mutants. (**A**) Single-plane spinning-disk confocal images of *S. pombe* of the indicated genotypes expressing the ER-luminal marker mCherry-ADEL and stained with BODIPY 493/503. (**A**-**B**) Loss of Dgk1 rescues the NE proliferation phenotype and the loss of nuclear circularity of *S. pombe spo7*Δ and *nem1*Δ mutants. Interestingly, the loss of Dgk1 does not rescue the large LD phenotype in the lipin mutants (**C**) but restores the average number of LD to the wild-type levels (**D**). (**E**) *dgk1*Δ cells with buckling spindle form unevenly sized daughter nuclei. Shown are the ratios of the larger to smaller daughter nuclei. (**F**) Representative time-lapse microscopy montage of mitotic *dgk1*Δ *S. pombe* that do not display obvious mitotic phenotypes (57% of cells). These cells show a similar decrease in NLS-DG sensor at the INM as compared to the wild-type (**G**). (**A**, **F**) Scale bars represent 5µm. (**B**-**D**) n=25 cells; (**E**) n=15 cells; (**G**) n=10 cells. p-values are derived from the unpaired t-test.

**Figure S5.**
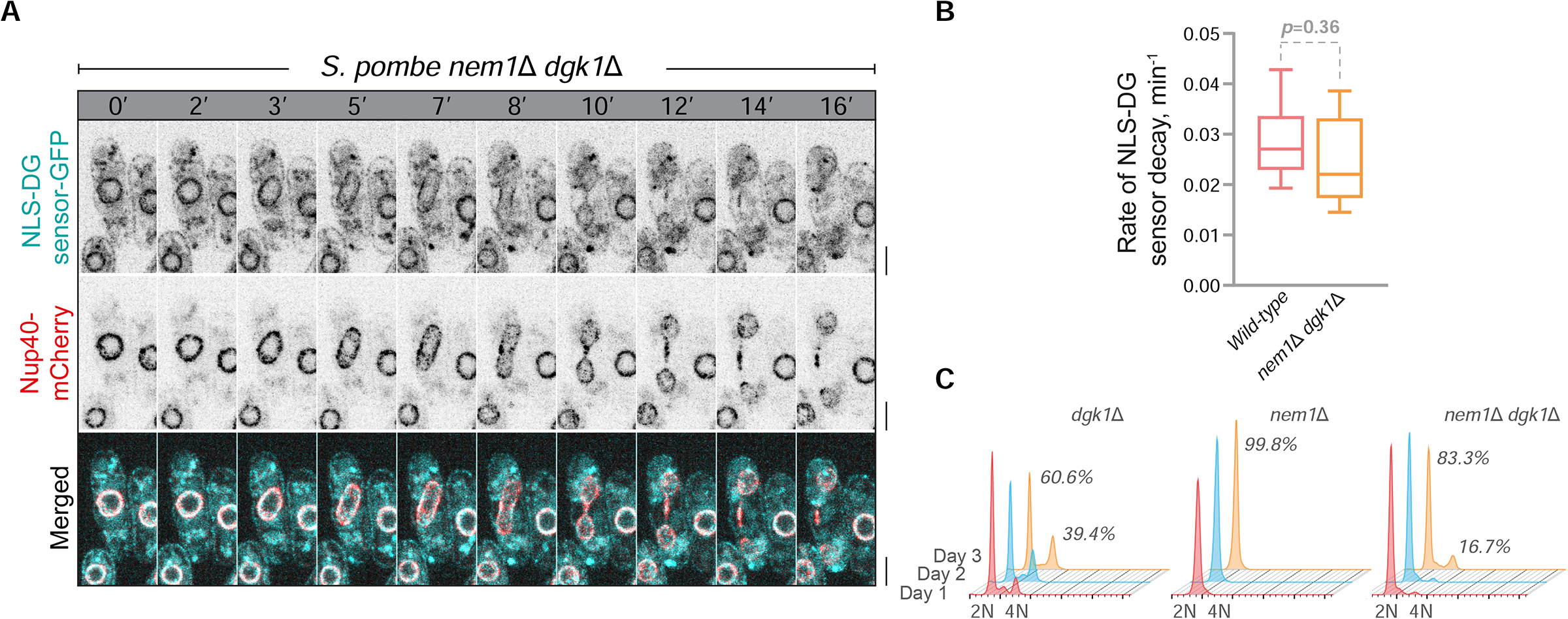
The mitotic phenotype of *S. pombe dgk1*Δ mutants can be rescued by the lack of the catalytic subunit of the Spo7-Nem1 complex, Nem1. (**A**) Time-lapse microscopy showing a similar decrease in NLS-DG sensor intensity at the INM in *nem1*Δ *dgk1*Δ double mutant as compared to the wild-type *S. pombe* during mitosis. Shown are the maximum projections of three Z-slices. Nucleoporin Nup40-mCherry marks the nuclear boundary; scale bars represent 5µm. (**B**) No significant difference in the rate of NLS-DG sensor decay is observed between the wild-type strain and *nem1*Δ *dgk1*Δ double mutant. (**C**) Deletion of *nem1* rescues the diploidization in *dgk1*Δ *S. pombe* as shown by flow cytometric analysis. Shown are the representative graphs from three biological repeats.

**Figure S6.**
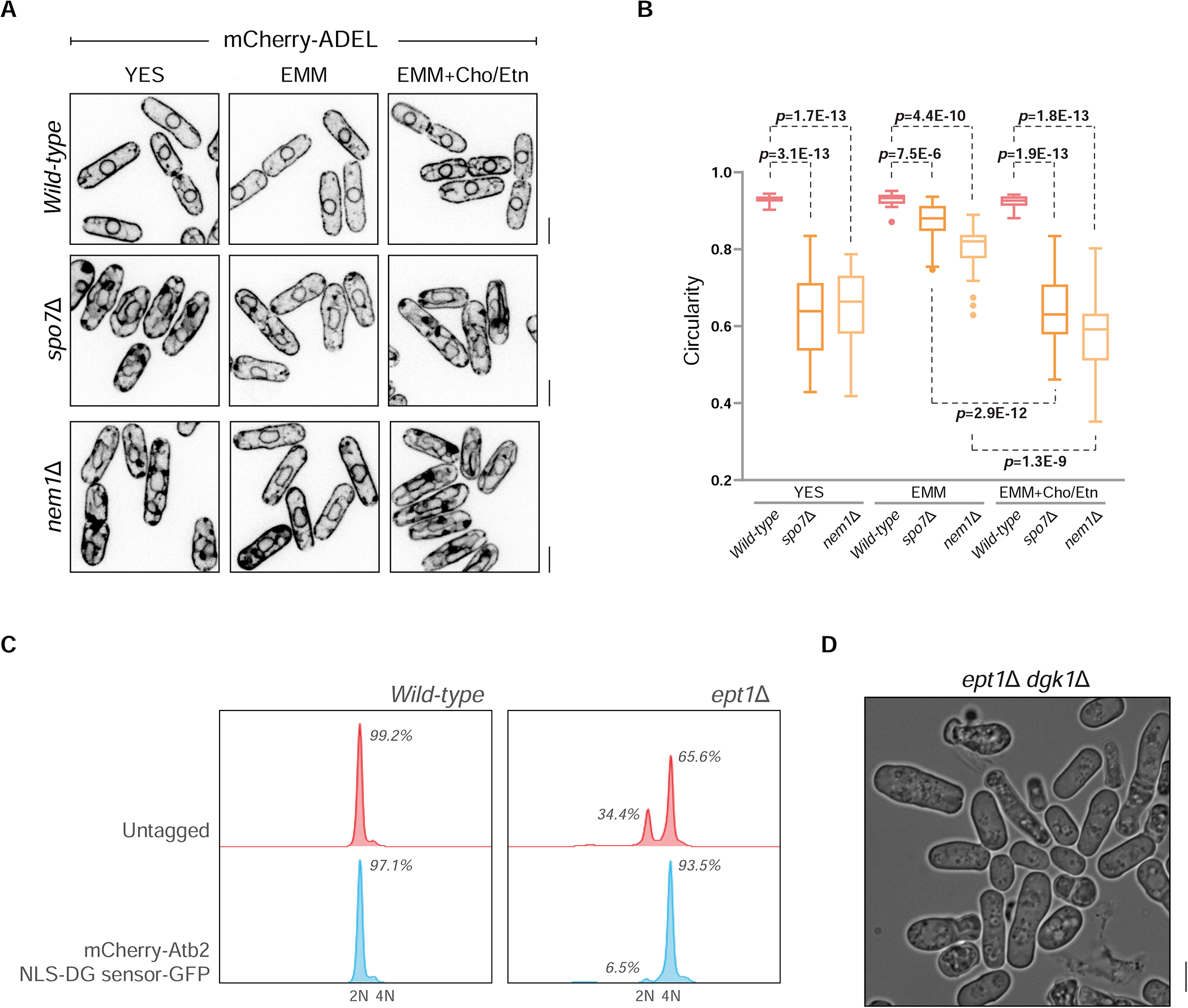
The Kennedy pathway strongly contributes to GPL biosynthesis in fission yeast. (**A**) Wild-type and lipin mutants expressing the ER-luminal marker mCherry-ADEL shows significantly less NE and ER expansion when cultured in the minimal EMM media. Supplementation of the Kennedy pathway precursors results in increased NE and ER expansion. Scale bars represent 5µm. Quantifications of nuclear circularity are shown in (**B**). n=25 cells. p-values are derived from the unpaired t-test. (**C**) Flow cytometric analysis shows that *S. pombe ept1*Δ mutants expressing α-tubulin mCherry-Atb2 are more susceptible to diploidization compared to the untagged strain, as shown for overnight cultures in rich YES medium. (**D**) High levels of cell death and large cells indicating the presence of diploids can be observed under phase contrast microscopy in the *ept1*Δ *dgk1*Δ double mutant strain.

